# Bioinformatics of combined nuclear and mitochondrial phylogenomics to define key nodes for the classification of Coleoptera

**DOI:** 10.1101/2024.10.26.620449

**Authors:** Thomas J. Creedy, Yinhuan Ding, Katherine M. Gregory, Luke Swaby, Feng Zhang, Alfried P. Vogler

## Abstract

Nuclear genome sequencing is resource-intensive and not practical for building densely sampled phylogenetic trees of the most species rich lineages of animals, while mitochondrial genomes can be sequenced and analysed with relative ease. Here, we develop a conceptual approach and bioinformatics workflow for combining nuclear single-copy orthologs with less informative but densely sampled mitochondrial genomes, for a detailed tree of Coleoptera (beetles). Basal relationships of Coleoptera were first inferred from >2,000 BUSCO loci mined from GenBank’s Short Read Archive for 119 exemplars of all major lineages under various substitution models and levels of matrix completion, to reveal universally supported nodes. Second, the corresponding mitogenomes were extracted and combined with an additional 373 species selected for broad taxonomic and biogeographic coverage, roughly in proportion to the known global species diversity of Coleoptera. Bioinformatic processing of mitogenomes was conducted with a novel pipeline for rapid, accurate annotation of protein-coding genes. Finally, phylogenetic trees from all 492 mitogenomes were generated under a backbone constraint from the universal basal nodes, which produced a well-supported tree of the major lineages at family and superfamily level. Being genetically unlinked and showing unique character variation, mitogenomes provide a unique perspective of the phylogeny. Comparison with three recent nuclear phylogenomic studies resulted in the recognition of >80 nodes universally present across all analyses. These may now support the higher classification of Coleoptera and serve as backbone of further studies, as numerous full mitogenomes and mitochondrial DNA barcodes are added to an increasingly complete phylogenetic tree of this super-diverse insect order.

## Introduction

Mitochondrial genomes remain an important source of data for phylogenetic reconstruction. They bridge a gap between disparate pursuits in taxonomic sequencing, in particular the ambitious plans for sequencing of full genomes for all species of eukaryotes (Lewin et al. 2022) and the equally ambitious endeavour of sequencing short mitochondrial barcode fragments for the study and monitoring of species across the world (Hobern and Hebert 2019). Mitochondrial genomes represent the phylogenetic ‘middle ground’ between complete genome assemblies and minimalist barcodes: they can be obtained for numerous species with relative ease, greatly surpassing nuclear genomes in taxon sampling and simplicity of analysis (Cameron 2014), while they contain the orthologous barcode region that can anchor barcode-based studies to the phylogeny (Li et al. 2023). Mitogenomes are widely used markers in evolutionary biology and population genetics (Desalle et al. 2017), and they are gaining new importance as a target of DNA sequencing from degraded museum specimens (Ferrari et al. 2023), as markers for species identification prior to genome sequencing (Blair 2023), and in large-scale characterisation of mixed biodiversity samples (Crampton-Platt et al. 2016). In addition, they are valuable as an independent data source for corroborating relationships inferred from nuclear genome data (Zhang et al. 2014)

Mitochondrial genomes have certain features that facilitate data processing and phylogenetic analysis. Across large portions of the Tree-of-Life, mitochondrial genomes exhibit a conserved structure, with a uniform number of genes, similar gene sizes, conservative gene order, and short intergenic distances of similar length (Cameron 2014). These features largely eliminate problems of ortholog assignment at the gene and nucleotide levels, allowing identification of the exact start and stop codons for each gene, and due to the lack of introns, to infer unambiguous reading frames from the primary sequence. However, mitochondrial genomes also are known for their high compositional and rate heterogeneity, producing long-branch artefacts from convergent biases in sequence variation (Cameron 2014). These issues can be partly ameliorated through the removal of the most strongly affected 3^rd^ codon positions (Hassanin 2006), the use of translated amino acid sequences (Rota-Stabelli et al. 2009), or the choice of advanced mixture models of sequence evolution that accommodate different evolutionary processes (Lartillot and Philippe 2004). Recent studies on tetrapods have shown that more complex models of sequence evolution that incorporate heterotachy and codon-specific sequence evolution improve the tree inferences, although true biological incongruence may partly be responsible for this “phylo-mito paradox” (Toups et al. 2024). As these effects of model violation affect the resolution of deep lineages most strongly, mitogenome trees may be combined with a scaffold of nuclear phylogenomic data to resolve the Tree-of-Life (Chesters 2017).

The number of available mitogenome sequences continues to increase rapidly (Cameron 2024), and the fast pace of data generation will need to be complemented with efficient methods for the bioinformatics of increasingly large datasets. While assembly of mitogenomes from shotgun sequencing is straightforward using a variety of strategies, software for genome annotations is lagging, as was pointed out in a recent review of insect mitogenomics (Cameron 2024). We address these challenges for the scaling-up of mitochondrial genomics and integration with nuclear phylogenomic data by devising a workflow that includes both data sources (Fig. 1). First, we develop a novel bioinformatics pipeline for semi-automated mitogenome data processing and error correction. This pipeline focuses on discriminating between ambiguous start/stop codons of the 13 protein-coding genes, followed by rapid alignment against a validated reference set of mitogenomes that cover the phylogenetic breadth of the target taxon. This novel set of scripts permits the automated polishing of mitogenome annotations after the primary annotation from established mitogenome processing pipelines. Second, we overcome the insufficiencies of mitogenomes in resolving deep lineages by combining them with nuclear phylogenomic data obtained from annotated genomes and shotgun short-read archives serving as a backbone for the mitogenome phylogeny. Vice versa, uncertainties in the topology from nuclear ortholog sets may be resolved by the mitogenomes due to their greater taxon sampling; thus, a critical step is to identify well supported nodes from the nuclear data while allowing the more densely sampled mitogenomes to contribute to the final resolution of basal nodes. Using this dual approach of combining (a few) nuclear and (many) mitochondrial genomes, the methodology generates a well-supported foundation for building a dense Tree-of-Life that may serve as a scaffold for integrating incoming new mitogenomes and for phylogenetic placement of rapidly growing DNA barcode and metabarcode data.

**Figure 1.**
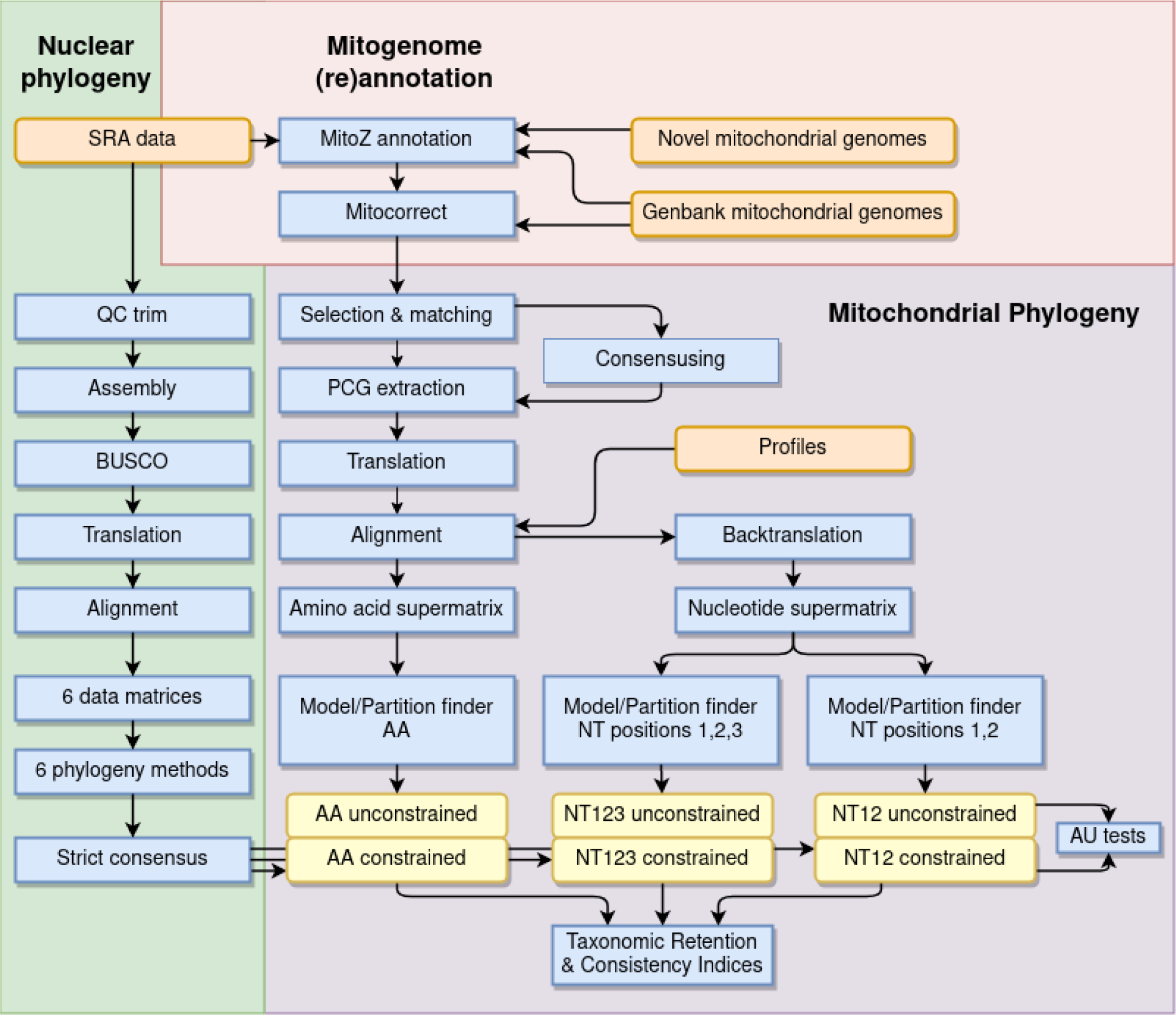
Workflow of data generation. Blue boxes indicate processes, orange boxes refer to input of data, and yellow refer to tree searches. Green section: Retrieving and processing nuclear phylogenomic data from SRA, followed by assembly (BUSCO), alignment, supermatrix generation and phylogeny under different model choice and tree search methods (see text). A strict consensus of trees across methods and models results in a single final nuclear tree, including unresolved nodes. Pink section: Generating the mitogenome data set by annotating (MitoZ) mitogenomes from SRA (or replacements, see text), unannotated mitogenomes from Genbank and novel mitogenomes, then improving (Mitocorrect) these annotations and those from annotated Genbank mitogenomes. Purple section: Mitogenome protein coding gene extraction, alignment and tree search from nucleotide and amino acid coding under various models, enforcing a backbone constraint based on the unequivocally supported nodes from the nuclear phylogenomic analysis.

We then employ this combined nuclear and mitogenome analysis to build an improved classification of the super-diverse and taxonomically challenging Coleoptera (beetles). Mitochondrial genomes have been useful for phylogenetics in Coleoptera at family and subfamily levels across the entire order, using the CAT model in PhyloBayes (Lartillot and Philippe 2004) to reduce long branch attraction problems (Timmermans et al. 2016). However, there are a few notable nodes that are not correctly resolvable even with advanced models, evident from clear contradictions with established taxonomies and nuclear genome trees. Two recent phylogenomic studies of Coleoptera using nuclear markers have been less affected by such systematic errors. Zhang et al. (2018b) used 99 PCR amplicons while McKenna et al. (2019) used ∼4,000 exons assembled from shotgun data for some 100 taxa, and when combined, these data covered most major lineages of Coleoptera at family level. These studies, and a recent re-analysis of the Zhang et al. (2018b) data by Cai et al. (2022), 2024), are broadly in agreement with each other and with the earlier mitogenome studies. However, poorly supported nodes and partly contradictory conclusions remain (Boudinot et al. 2022), and no formal comparisons between the nuclear and mitochondrial datasets have been made. We generated a taxonomically and biogeographically comprehensive mitogenome set for some 140 families of Coleoptera and integrate these data with mined nuclear orthologs, benefitting from the combined power of greater taxon density of mitogenomes and the greater reliability of nuclear genome data at the deep level, to resolve a set of universally supported nodes that define the basal relationships of Coleoptera. Our workflow also includes an algorithmic method for comparison of nuclear and mitochondrial derived trees, which usually have been based on qualitative assessments only (Cameron 2024).

## Materials and Methods

### Building the backbone from nuclear genome sequences

The GenBank Sequence Read Archive (SRA) was searched using the web-based portal for raw genome and transcriptome sequence data for Coleoptera, filtering for libraries with >5 Gb of data that generally corresponds to ∼20x coverage for most insect genomes required to obtain a largely complete ortholog sets. We selected 110 taxa that represented the greatest coverage available at the family level, and added 9 sequences from the Strepsiptera, Megaloptera, Neuroptera and Raphidioptera as outgroups. Quality trimming of raw data was performed with fastp (Chen et al. 2018). The read data were used for transcriptome assemblies with SOAPdenovo-Trans (Xie et al. 2014) and genome assemblies with SOAPdenovo2 (Luo et al. 2012), respectively. Duplicated transcripts and artefacts were reduced by CD-HIT (Fu et al. 2012). To extract single-copy orthologs from the assemblies and to assess the completeness of assemblies, we applied Benchmarking Universal Single-Copy Orthologs (BUSCO) analyses using the Endopterygota set (Waterhouse et al. 2019). The parameter configurations of most gene prediction tools need to be optimised on each specific genome, but BUSCO can enhance genome annotation procedures by providing high-quality gene model training data even in the absence of support from orthologs or transcriptome evidence of closely related species. BUSCO can identify matching single-copy orthologs in a given taxonomic group from many sequenced and annotated genomes of the major species clades based on the OrthoDB catalogue of orthologs. It allows gene duplications or losses during evolution, and the filtering of single-copy genes is controlled by thresholds, while the incompleteness of available assembled genomes and predicted gene sets is also taken into account (Waterhouse et al. 2018; Manni et al. 2021). Sequence alignment on each orthology gene set was conducted with Mafft v7.450 (Katoh and Standley 2013), followed by alignment trimming with BMGE v1.12 (Criscuolo and Gribaldo 2010) based on the BLOSUM90 substitution matrix, and concatenation of the filtered orthologs for final matrix generation with FASConCAT-g v1.04 (Kück and Longo 2014).

To reduce possible systematic error, we conducted phylogenetic tree searches using a diverse set of analytical methods. First, we generated six data matrices by concatenating loci according to two criteria of completeness: (i) loci that are present in a minimum percentage of taxa (equal to or greater than 50%, 75% or 90%) and (ii) loci that have a minimum proportion of gene sequence present in the taxa where they are present (greater than 0% or 70%). All matrices were translated into amino acid format. Tree searches were conducted using maximum likelihood (ML) in IQ-TREE v2.0-rc2 (Minh et al. 2021) under various substitution models, including: (1) the Dayhoff fixed replacement rate matrix (Dayhoff 1978); (2) the improved LG generalised replacement rate matrix that incorporates evolutionary variability of rates across sites (Le and Gascuel 2008) (IQ-TREE options: -m MFP --mset LG --msub nuclear --rclusterf 10); (3) the EX_EHO site-specific mixture model which estimates replacement matrices separately for sites based on six different secondary structure category partitions (Le and Gascuel 2010) (IQ-TREE options: -m MFP –mset EX_EHO); (4) a site-specific mixture model, based on stationary exchange probabilities for up to 60 separate profiles whose codon frequencies may be estimated from the data (Le et al. 2008), implemented as the posterior mean site frequency (PMSF) model using the LG tree as an initial guide tree (Wang et al. 2018) (IQ-TREE options: -m LG+C60+F+G); (5) the edge-unlinked GHOST (General Heterogeneous evolution On a Single Topology) mixture model that accounts for rate variation across sites and across lineages (heterotachy) (Crotty et al. 2020) (IQ-TREE options: -m LG+F0+H4). In all IQ-TREE phylogenies, node support values were calculated using 1,000 SH-aLRT replicates (Guindon et al. 2010) and 1,000 ultrafast bootstraps (Hoang et al. 2018). For the LG and EX_EHO models we employed IQ-TREE Partition/ModelFinder (Kalyaanamoorthy et al. 2017) to implement partition models (Chernomor et al. 2016). A sixth approach employing the multi-species coalescent model (MSCM) addressing Incomplete Lineage Sorting (ILS) was implemented using ASTRAL-III v5.6.1 (Zhang et al. 2018a) based on the set of gene trees estimated with IQ-TREE to infer coalescent-based species trees, with local branch support estimated from quartet frequencies (Sayyari and Mirarab 2016). We quantified genealogical concordance with the gene concordance factor (gCF) and the site concordance factor (sCF) given the reference tree and gene trees (Minh et al. 2021) using IQ-TREE.

To capture the topological variation among the 36 analyses (six matrices by six models), we generated a strict consensus of trees from each combination of matrices and models. We then computed the Robinson-Foulds (RF) (Robinson and Foulds 1981) distance matrix between all consensus trees and ordinated this matrix using Non-metric MultiDimensional Scaling (NMDS). This provided a measure of similarity among the tree topologies obtained from a specific data matrix under different models. We assessed these trees for their congruence with the Linnaean taxonomy at three hierarchical levels (suborder, superfamily, family) using the taxonomic Retention Index (tRI) (Hunt and Vogler 2008) to identify the tree that best agreed with the existing classification of Coleoptera. This approach treats presence or absence in each taxonomic group as a character state, computing the classic Retention Index (RI) for each taxon that is then averaged across all taxa for the particular hierarchical level to give a single value. Finally, a strict consensus tree was computed from individual consensus trees to obtain a set of nodes that was robust to both the model choice and the completeness of the data matrix; this tree formed the backbone for subsequent mitochondrial phylogenetics.

### Mitogenome acquisition and annotation

We generated a database of Coleoptera mitogenome sequences, sourced from Genbank and novel sequencing and assembly. Retrieval from Genbank took place in April 2023 for this study, using a web portal search (search term “Coleoptera” “mitochondrial” “mitochondrion sequence” AND “genomes” NOT “predicted” NOT “shotgun”), restricting retrieved sequences 3-50 kb in length. The result of this search was combined with thousands of unpublished mitogenomes in our database (Creedy, Lee et al., unpublished). This set was used to build a preliminary exploratory tree aiming to maximise representation at the family and subfamily level, generally by selecting two distantly related representatives of a family or representatives of multiple subfamilies. Mitochondrial genomes with a complete set of protein coding sequences were used preferentially. We also selected for geographic breadth, to ensure a worldwide representation on the final phylogeny, including a few unpublished sequences from tropical sites. Our final selection comprised 373 mitochondrial genomes.

Next, the joint analysis with the nuclear data required that all species with nuclear genomes were represented by the corresponding mitogenome. For 80 of the 119 terminals in the nuclear phylogeny the assemblies of SRA data produced mitochondrial genome contigs that were complete for the 13 protein coding genes. For the remaining 39 terminals either no mitochondrial contig was assembled, the assembled genome was incomplete, or it was of very poor quality, perhaps due to varying stages of post-transcriptomic processing. For each of these 39 species, we searched our mitogenome database for close relatives. In 33 cases, the closest relative available in the database was a substantially more complete conspecific (31) or congener (2), and we simply substituted our database mitogenome for the missing, incomplete, or poor SRA assembly. In five cases, the SRA assembly was incomplete, an incomplete conspecific was available in the database, but the combination of both sequences would increase the number of Protein Coding Genes (PCGs). In these cases, we aligned and generated consensus sequences for each PCG, using ambiguity codes for any polymorphisms. In the final case, *Haliplus*, the SRA assembly was very poor but the closest relative in our database was an incomplete congener. Given the importance of this representative for the tree and the poor availability of mitogenomes for this taxon, we used a consensus of these sequences in this case. We thus successfully identified suitable mitochondrial sequences to correspond with all 119 nuclear terminals, for a final count of 491 mitogenomes.

All mitochondrial genomes (SRA assemblies, unpublished sequences, GenBank sequences) were annotated for PCGs, RNA genes and tRNAs using the MitoZ pipeline (Meng et al. 2019). The PCG annotations of all mitochondrial genomes were refined using the newly developed python tool *mitocorrect* (https://github.com/tjcreedy/mitocorrect) (Fig. 2). The script evaluates alternative potential start and stop codons based on two axioms: (i) given the conserved structure of mitogenomes in Coleoptera, the distance (or short overlap) between adjacent genes generally differs only by a few base pairs around a modal number; (ii) presuming conserved lengths of the protein sequences, alignments of each PCG can indicate the most probable start and stop codons relative to other sequences in a multiple sequence alignment, from which any new annotation may deviate by a certain number of bases. *mitocorrect* evaluates each potential start and stop codon, balancing the two criteria for an in-frame nucleotide sequence (Fig. 2). The parameter values for the range of gene distances and admissible start/stop codons for each gene are provided by the user and based upon well-established understanding of mitogenome PCG properties regarding the modal distances of genes and homologies of start and stop codons provided to the program in a specifications table (Supplementary text S1).

**Figure 2.**
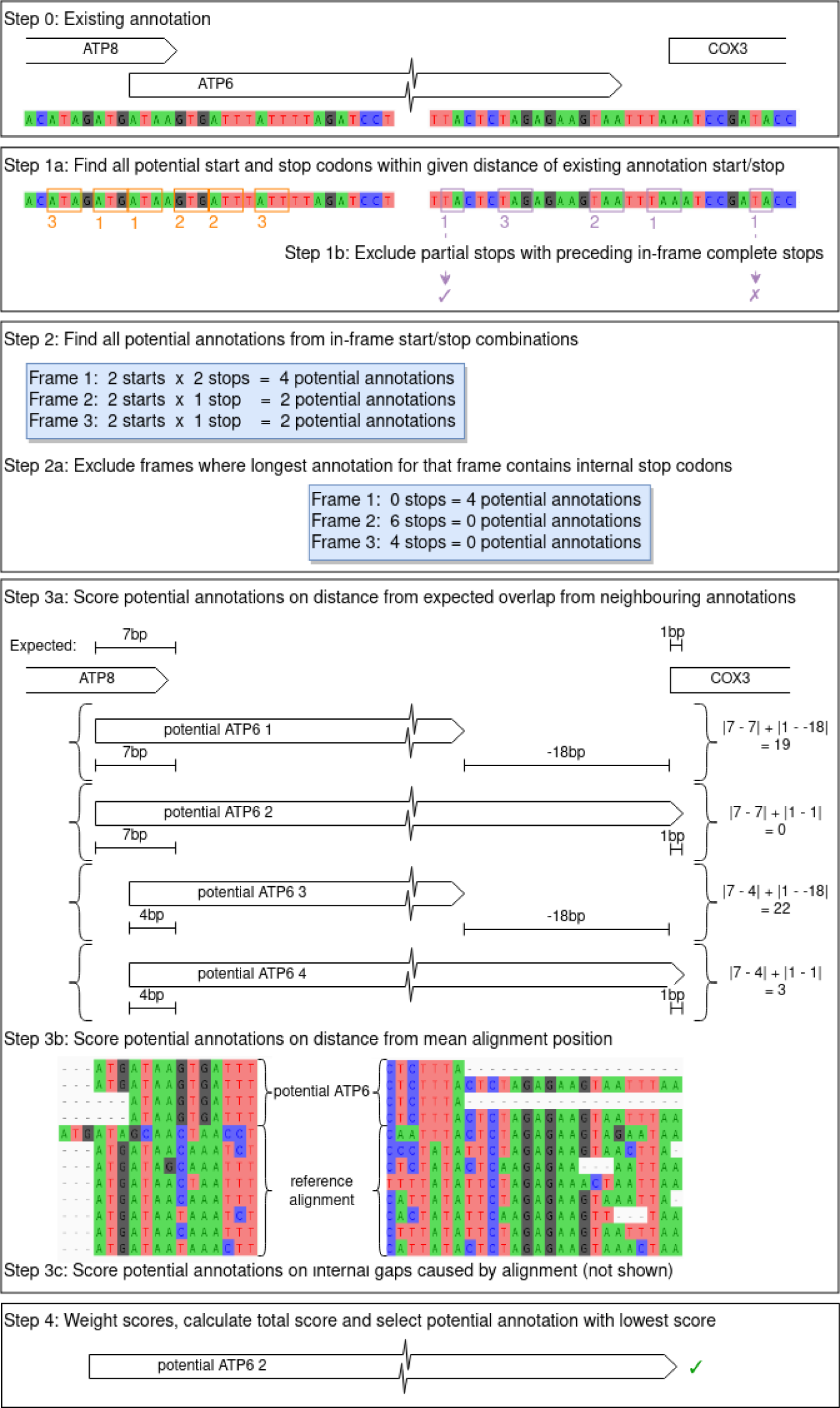
The mitocorrect methodology. Alignment and topography together determine the selected start and stop codons. In the example, the starting annotation’s start and stop positions are valid codons, but these may not be correct (step 0). Multiple alternatives are available (step 1) but many of these are rejected for being in the incorrect frame (step 2), including the initial stop position. All possible remaining combinations form the potential annotations (step 2a). Comparison with the expected gene overlaps or gaps, supplied by the user based on external information, allows scoring of potential annotations (step 3a). Further evidence is generated through comparison of the potential annotations to a reference alignment (step 3b and c). These comparisons are scored for their deviation from the modal gene distance and alignment matches (in number of nucleotides), and the overall lowest scoring annotation is selected.

### Mitochondrial gene alignment

The new annotations were used to excise the 13 PCGs of the 491 input mitogenomes and each gene was aligned based on the amino acid translation. First, nucleotide sequences were translated to amino acid sequences, then these were aligned using the MAFFT (Katoh and Standley 2013) *ginsi* global pairwise alignment strategy. The amino acid alignments were translated back to nucleotide sequences by taking the unaligned sequences and inserting gaps in the appropriate positions as determined by the amino acid alignment, implemented in a python script (https://github.com/tjcreedy/biotools/backtranslate.py). The two sets of 13 alignments (nucleotide and amino acid) were converted into two supermatrices using catfasta2phyml.pl (Nylander 2010). Ribosomal RNA genes were not included in the analysis.

### Mitochondrial phylogenetic analysis

For the aligned data, we considered three possible datasets for phylogenetic reconstruction: (i) the complete nucleotide supermatrix with all nucleotides included (NT123), (ii) the nucleotide supermatrix excluding 3^rd^ codon positions (NT12), and (iii) the amino acid supermatrix (AA). For each dataset, we started from a maximal partition scheme, namely (i) gene and codon position partitioning (39 partitions), (ii) gene and first two codon position partitioning (26 partitions), and (iii) gene partitioning (13 partitions) respectively. We then used the combined partition and model finding algorithm of IQ-TREE (v2.0.3) to first implement a PartitionFinder-like greedy partition merging approach to find the preferred partition scheme (Lanfear et al. 2012), then applied automatic model selection (Kalyaanamoorthy et al. 2017) to find the most appropriate model for each partition. This outputs a partition and model table for each of the three datasets.

For each dataset, we used IQ-TREE (v2.0.3) to reconstruct the ML tree based on the input dataset and partition/model table. To assess branch support, we used the ultrafast bootstrap (Hoang et al. 2018) with 1,000 replicates. We also performed three single branch tests to provide further support values: the SH-like approximate likelihood ratio test (Guindon et al. 2010), the approximate Bayes test (Anisimova et al. 2011) and the fast local bootstrap probability method (1,000 replicates) (Adachi and Hasegawa 1996). These analyses were carried out separately (i) without constraint and (ii) with a multifurcating backbone constraint phylogeny, representing the consensus of the various nuclear tree topologies obtained as described above.

### Comparison of mitochondrial phylogenies to published trees

To explore consistency in node recovery among recent studies of the Coleoptera, we acquired or transcribed the topology of the trees produced by Zhang et al. (2018, Figure 2), McKenna at al. (2019, Figure S1), and Cai et al. (2022). To ensure consistent taxonomy among trees, we matched terminals by the reported species, genus and/or family in these trees to the taxonomy of our mitogenome database, then for any terminals still without taxonomy, we retrieved taxonomy from NCBI. Based on our knowledge of critical nodes in the phylogeny of Coleoptera and comparisons of our mitogenome based tree with the above studies, we identified a total of 84 clades comprising one or more taxa at family or higher levels recovered consistently with the potential for defining a stable higher-level classification of the Coleoptera. For each phylogeny, we identified the set of terminals that belonged to each of these clades, which we refer to as a “taxonomic set”; we then located the most recent common ancestor (MRCA) node for each taxonomic set. Based on the subtree below this node, we classified each taxonomic set on each phylogeny as:

1. *Monophyletic*: all terminals corresponding to a given taxonomic set *present in the tree* are descended from this node.
2. *Monophyletic-incomplete*: all terminals corresponding to a given taxonomic set present in the tree are descended from this node, but other taxa in this tree *not assigned to any other taxonomic set* are also descended from this node; this category is relevant mainly to families (or groups at other levels) not consistently sampled across various studies and thus not assigned to any taxonomic set.
3. *Paraphyletic*: all terminals corresponding to a given taxonomic set present in the tree are descended from this node, but members of other taxonomic sets are also descended from this node, and thus *this node may also be the MRCA of another taxonomic set*.
4. *Absent*: the taxonomic set is not represented in the tree or is represented by only one exemplar.

This analysis was carried out using the *taxize* 0.9.98 (Chamberlain and Szöcs 2013), *ape* 5.6.2 (Paradis and Schliep 2019) and *phangorn* 2.10.0 (Schliep 2011) packages in R 4.2.2 (Team 2013), along with custom functions.

### Compositional heterogeneity

Compositional heterogeneity was explored using the Relative Compositional Frequency Variation (RCFV) (Zhong et al. 2011). The metric uses the relative frequency of a nucleotide or an amino acid in a given taxon against the mean relative frequency of the same nucleotide or amino acid over the entire dataset. The analyses were performed for the nucleotide dataset with all three codon positions included (NT123), after removing the third codon position (NT12), and on the amino acid dataset (AA). Within each dataset, each partition derived from the partition merging approach (see below) was considered independently. Analyses were conducted using a custom R functions *rcfv* and *rcfv.partitioned* (https://github.com/tjcreedy/phylostuff/blob/main/phylofuncs.R). For each dataset and partition, we then graphically explored the partitioning of overall RCFV into character-specific (csRCFV) and taxon-specific (tsRCFV) components (Fleming and Struck 2023) and used chi-squared tests to test whether the distribution of csRCFV and tsRCFV values across character sets and taxa respectively deviated from random chance, i.e. to test whether any characters or taxa appeared to have significantly greater compositional frequency and thus mislead the phylogenetic reconstruction.

## Results

### Nuclear gene assemblies

Successful assemblies of nuclear orthologs were achieved for 119 diverse taxa of Coleoptera and outgroups (Supplementary Table S3), selected to cover the greatest phylogenetic diversity available at the SRA (September 2020). Several key taxa for which shotgun data have been reported in the literature were missing from the SRA and thus could not be included in the current study. The final data matrix extended to 527,095 amino acids, representing 2,127 independently assembled loci (Table 1). The number of genes present in each terminal dropped off sharply under more stringent requirements for matrix completion (moving from ≥50% to ≥75% of genes present) or sequence completeness (≥70% of nucleotides present) (Table 1). In total, we obtained sequences for 78 families covering the four suborders of Coleoptera and all 16 widely recognised superfamilies of the large suborder Polyphaga. Taxon sampling within Adephaga included all known family-level taxa, except for Meruidae, which is closely related to Noteridae. For the two small suborders, Myxophaga and Archostemata, representatives of two families each were assembled successfully.

**Table 1:**
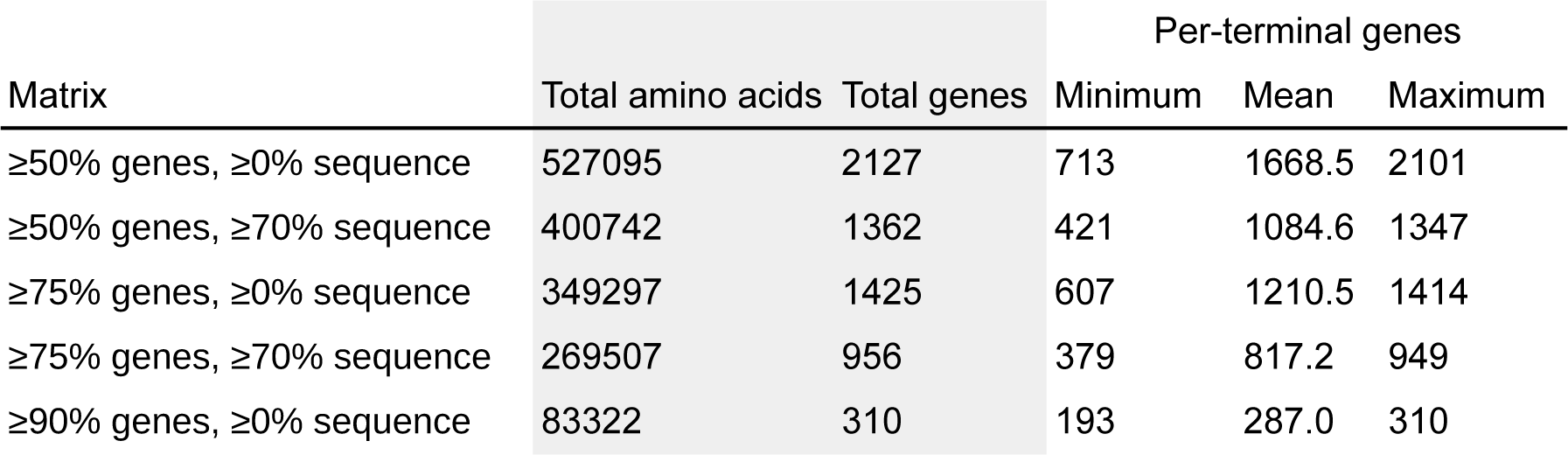
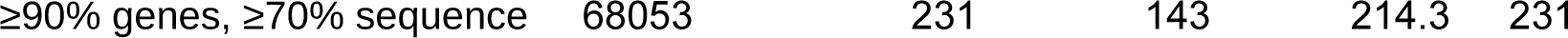
Summary of Nuclear Datasets. Six datasets fulfilling differing levels of two completeness criteria were generated. For each dataset, the number of amino acids and genes in the resulting supermatrix is reported. The mean and range of the number of genes per terminal in the supermatrix is also given. Finally, the table shows for the two approaches where PartitionFinder was used, the final number of partitions with different models/parameters in the tree search. All 119 terminals were present in all matrices.

Phylogenetic trees based on data matrices of various levels of completeness and generated under different substitution models (36 trees in total, Supplementary Material S4) were ordinated based on a matrix of all pairwise RF distances (Fig. 3). All topologies from the 50% and 75% among-gene completeness matrices were similar under any of the models, except for the trees generated with ASTRAL that occupied a small portion of the multivariate space distant from all others (Fig. 3). Only the topologies from the 90% completeness matrices were widely different from all others, and from each other, showing that this most complete but smallest matrix remained sensitive to the model choice, unlike the larger matrices (Fig. 3).

**Figure 3.**
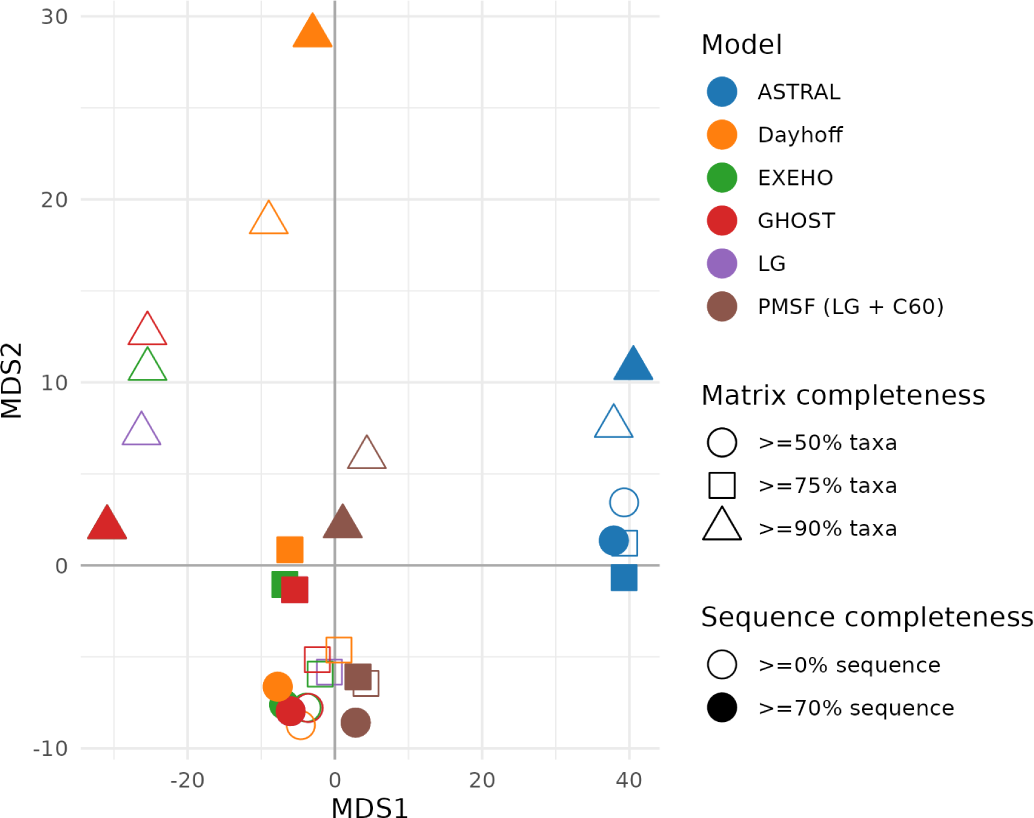
Ordination of RF distances of trees from nuclear genomic data using different matrix types and substitution models. Symbols correspond to six models, each of which were used to generate a tree under the six different matrices (3 levels of across-gene completeness, colours, 2 levels of within-gene completeness, filled vs unfilled). Where more than one best tree was obtained in a search, the RF distances were calculated from the consensus tree. The tree from the matrix with 90% matrix completion and 70% gene completion obtained under the PSMF model (green triangle at centre of graph) was used to illustrate the consensus among tree topologies in Fig. 4. Note that the trees returned for LG and GHOST models were frequently identical or highly similar for several matrix completeness cases, so these LG (purple) points are obscured by the GHOST (red) points in these cases.

Phylogenetic relationships encountered across the 36 analyses were captured well in a tree from the 90% matrix completeness dataset under the PMSF model, which was at the centre of the multivariate space. Against this exemplar topology (Fig. 4), we scored the nodes that were found in all of the other analyses (i.e. all nodes in this tree based on 310 genes were also present in trees of up to 2,127 genes but with lower matrix completeness; Table 2), as a measure of sensitivity to the matrix type and model. We first observed that the ASTRAL analysis added a great number of unique nodes, including the non-monophyly of Coleoptera, and therefore this method was removed from further analysis. For the remaining 30 trees, nodes sensitive to matrix choice and model were clustered in certain parts of the tree, especially in Tenebrionoidea and Scarabaeoidea. Several nodes were consistently found to be resolved differently under various parameter sets (matrices and models), while several others were affected only by one parameter set (shown as a heat map on the nodes of Fig. 4). Especially, the topology obtained with the Dayhoff replacement matrix differed in several basal nodes, including the node grouping the two suborders Archostemata and Myxophaga, and the node grouping Cleroidea and Coccinelloidea (Fig. 4). Generally, we found that if a node was not present across multiple data matrices it was also not present across multiple models within one type of matrix, i.e. inconsistent nodal support due to limited matrix completion coincides with sensitivity to the model. In sum, 83 nodes were resolved universally across these trees under a strict consensus (Supplementary Material S5), which were then used as the backbone constraint for mitochondrial tree searches (see below).

**Figure 4.**
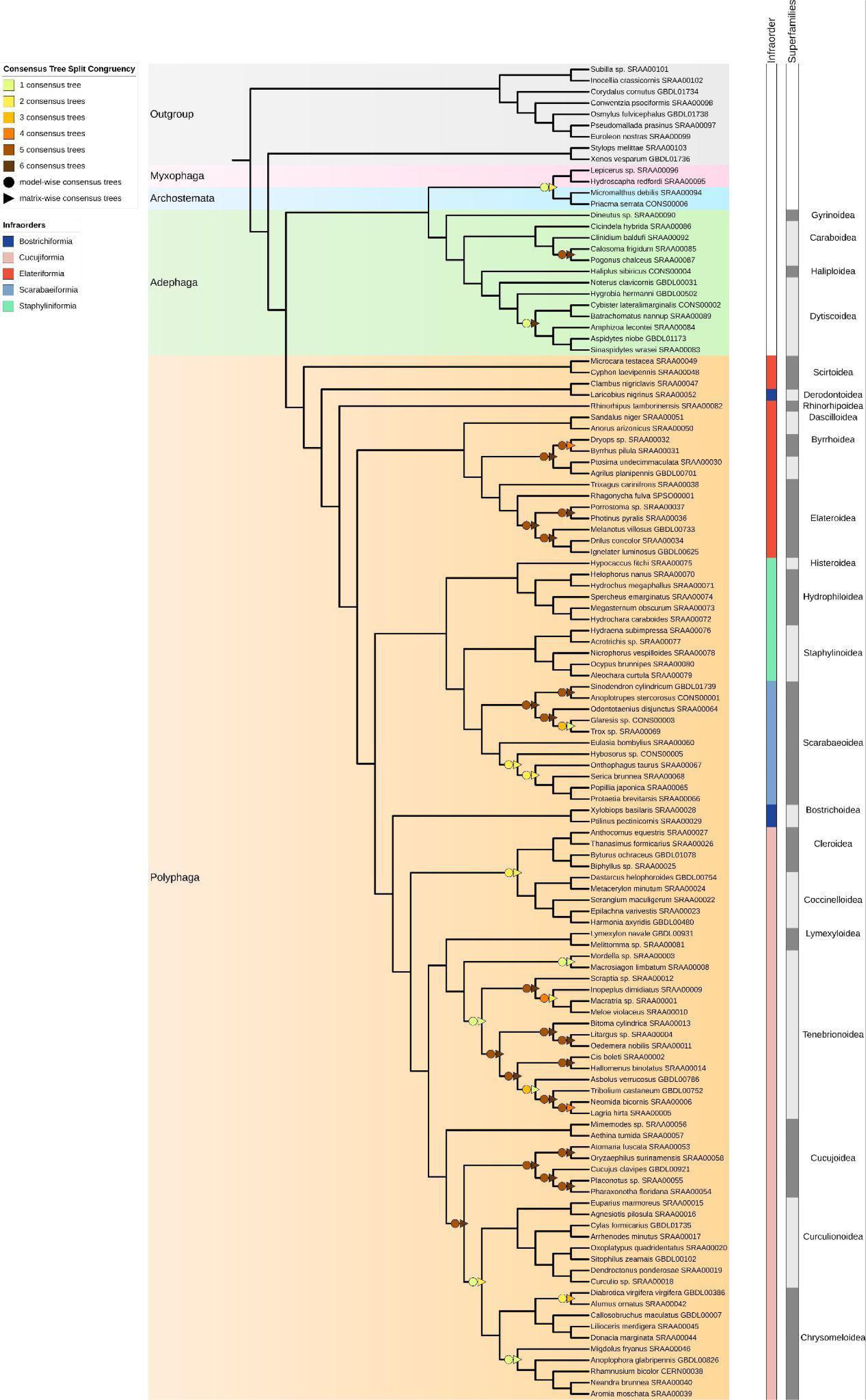
Exemplar tree of Coleoptera from phylogenomic data. The tree is based on the matrix_90 dataset and the PMSF model. The symbols indicate the nodes that are not obtained on the trees from other data matrices (triangles) and models (circles), and the colour indicates how many of the five models (excluding ASTRAL) or six matrices do not include these nodes. Infraorders are based on the NCBI taxonomy.

**Table 2.**
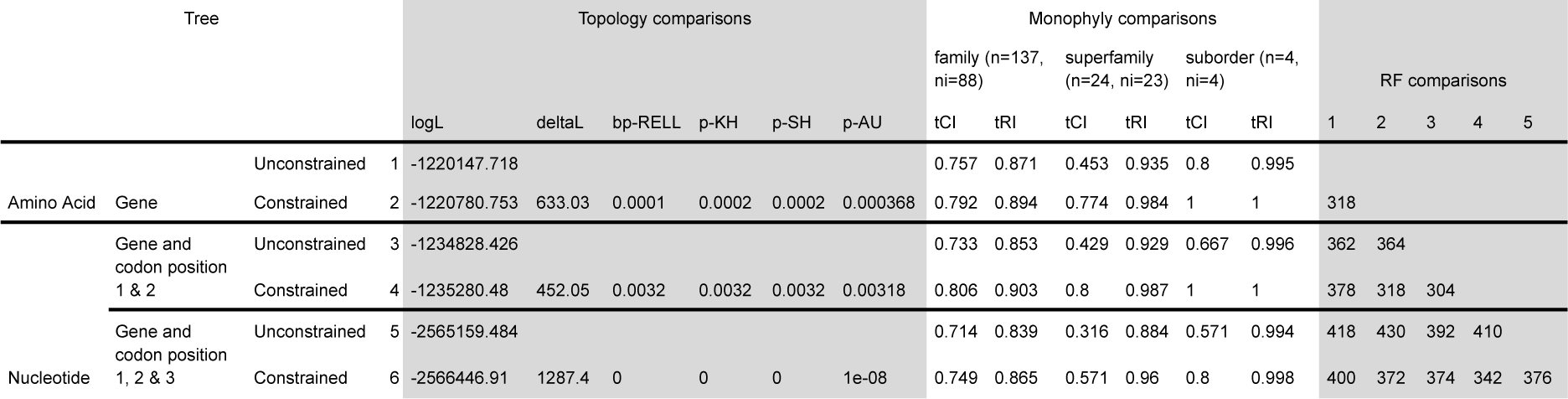
Tree topological tests and monophyly tests comparing the backbone constrained versus unconstrained tree searches. Topology comparisons gives the probability that a tree is significantly different from the best tree (outside of the 95% confidence interval), for a variety of tests including Resampling of Estimated Log Likelihoods (bp-RELL), Kaneshiro-Hasegawa (KH), Shimodeira-Hasegawa (SH), and Approximate Unbiased (AU) tests. For monophyly comparisons, the number of unique taxa (n) and the number of informative taxa (ni, the number of taxa with >1 tips) is given for each taxonomic level tested. tCI and tRI refer to the ensemble (mean) taxonomic consistency and retention indices, respectively. RF comparisons present pairwise RF distances between trees according to the numbers assigned to each row.

### Mitochondrial data matrices

Mitochondrial genomes were assembled from the shotgun database and available mitogenomes at Genbank, supplemented with a few unpublished mitogenomes to maximise the taxonomic diversity at the family and subfamily levels and to represent the highly uneven species richness of lineages of Coleoptera. We aimed at one mitogenome per ∼1,000 species for larger families but also included phylogenetically unique, species-poor lineages. Assembly of mitochondrial genomes from transcriptome data were frequently poor and were substituted with available mitogenome sequences for the same species or a close relative (see Material and Methods, and Supplementary Materials S3). The final set of 482 ingroup mitogenomes represented the major taxonomic and geographic lineages of the Coleoptera, including representatives of 138 families and 176 subfamilies (Supplementary Materials S6), with a matrix completion at gene level of 87%. Where necessary the 5’ and 3’ ends of each gene annotation were adjusted using *mitocorrect*, producing the best estimate of start and stop codons for the final alignment procedure. Across all mitogenomes, the gene organisation was highly conserved. Generally, adjacent genes either overlapped by one or a few nucleotides, or they were separated by a small number of intergenic positions. Only in rare cases were these distances >100 nucleotides or more, especially in the region between the *nad1* and *cytb* genes that are transcribed in opposite directions. Presumed start codons were highly variable, with up to nine different start codons in *nad2* and *cox1*. Due to the conservative gene distances (and conservative gene order) the *mitocorrect* procedure could be limited to searching a range (“search distance”) of 50 nucleotides or less from an existing annotation (obtained from Genbank or using standard annotation software, e.g. MitoZ), which limited the number of potential start/stop codon combinations to be tested. Mitochondrial PCGs were excised from the annotated mitogenomes and aligned as translated amino acid sequences before being back-translated for nucleotide-based analysis (Materials and Methods). The final matrices of the 13 PCGs included 12,270 nucleotide positions or 4,090 amino acid positions (Supplementary Material S7).

### Tree searches and topology

Tree searches were conducted on three matrices of DNA (all nucleotide sites; 1^st^ and 2^nd^ codon positions only; amino acid transformed data) with and without the constraint (Supplementary Materials S8). Phylogenetic trees generated without constraint produced topologies in broad agreement with known mitogenome trees and had overall good fit with the Linnaean taxonomy at family level and above based on the tRI (Table 1, Fig. 5). However, these trees were also marred by a few severely misplaced groups and inconsistent relationships of deeply branching lineages, in particular for the trees that did not exclude the 3^rd^ codon positions. Applying the backbone constraints from the 83 universally resolved nodes of the nuclear gene trees produced topologies of lower likelihood than the unconstrained trees at a level that was significant in all commonly used tests of tree selection (Table 1). However, the tRI was higher at all hierarchical levels indicating a better fit to the Linnaean classification, although even the constraint searches did not avoid a few nonsensical groupings that mostly consisted of clusters of species with no close relatives in the nuclear dataset constraint. Using the amino acid sequence improved the topology, but a few such cases remained, including the three representatives of the Osoriinae (Staphylinidae) and two species of Sphindidae placed very distantly from their likely positions (which were ignored in the subsequent analyses).

**Figure 5.**
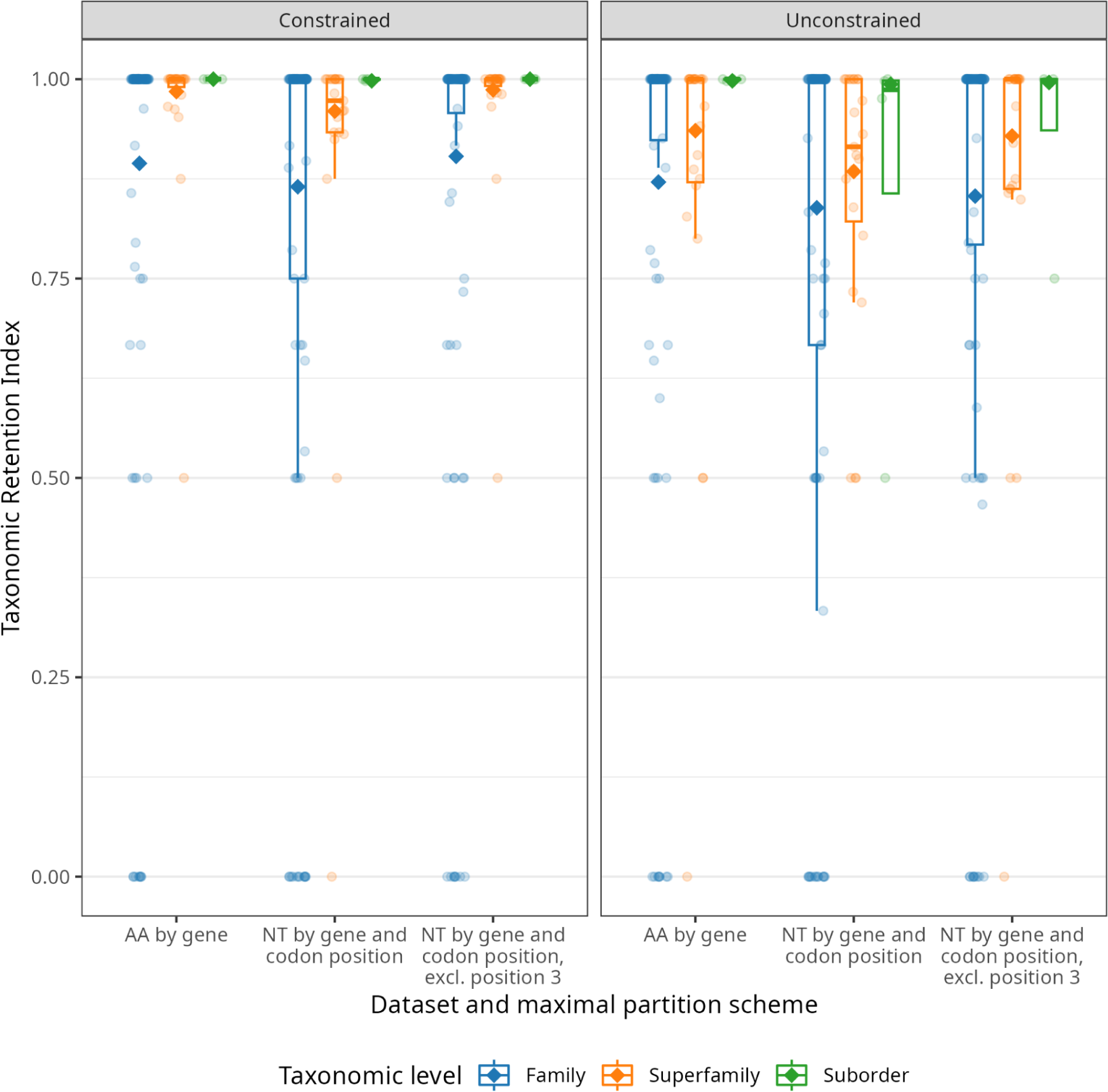
Match of mitochondrial phylogeny and Linnaean taxonomy. The taxonomic Retention Index (tRI) for individual taxonomic groups (circular points and boxplots) at three hierarchical levels (colours) with (left panel) and without (right panel) enforcing the backbone tree. Filled diamonds show the consensus tRI across all taxonomic groups for a given taxonomic level and tree. A small amount of x-axis jitter has been applied to circular points for improved clarity.

Using the RCFV metric for assessing compositional heterogeneity demonstrated that the poorly performing partitions indeed showed greater heterogeneity. Specifically, the partitions P3 and P6 exclusively composed of 3rd codon positions showed the highest levels of heterogeneity, followed by the partition P4 and then by P1 and P5 all of which are predominantly composed of 1st codon positions. Partition P2 composed of 2nd codon positions had the lowest RCFV values and showed limited variation in these values (Supplementary Figure S7). Character-specific compositional heterogeneity (csRCFV) showed similar levels of heterogeneity for the four nucleotides, although this increased for A and T in the 3rd codon positions (partitions P3 and P6). The taxon-specific composition heterogeneity (tsRCFV) revealed levels of heterogeneity largely in line with the overall RCFV values for each partition, but generally with a long tail of high values, especially in P2 of the amino acid analysis, which otherwise had lowest overall heterogeneity. Cases of high tsRCFV may indicate the underlying causes of severely misplaced taxa, but we did not find this correlation when mapping tsRCFV values on the phylogenetic tree.

### Classification of Coleoptera

The tree generated from the amino acid data under the backbone constraint provided a detailed perspective on family level relationships of Coleoptera (Fig. 6). Discussing briefly the major groups we noted: (i) The relationships of the four suborders have been problematic, but nuclear genome data (Misof et al. 2014) resolved them as (Outgroup (Polyphaga (Adephaga (Myxophaga, Archostemata))). This topology was also supported in all our phylogenomic analysis, except for the Dayhoff rate model with the 90% complete dataset (*matrix_90 abs70*), which supported the sister relationships of Adephaga and Archostemata, albeit with low support. Inclusion of this tree in the selection of the backbone constraint also removed the node that specified the suborder relationships from the backbone, and thus the mitogenome signal was tested here, which grouped Adephaga and Myxophaga, as generally seen in mitogenome data except with the superior CAT model in PhyloBayes (Timmermans et al. 2016).

**Figure 6.**
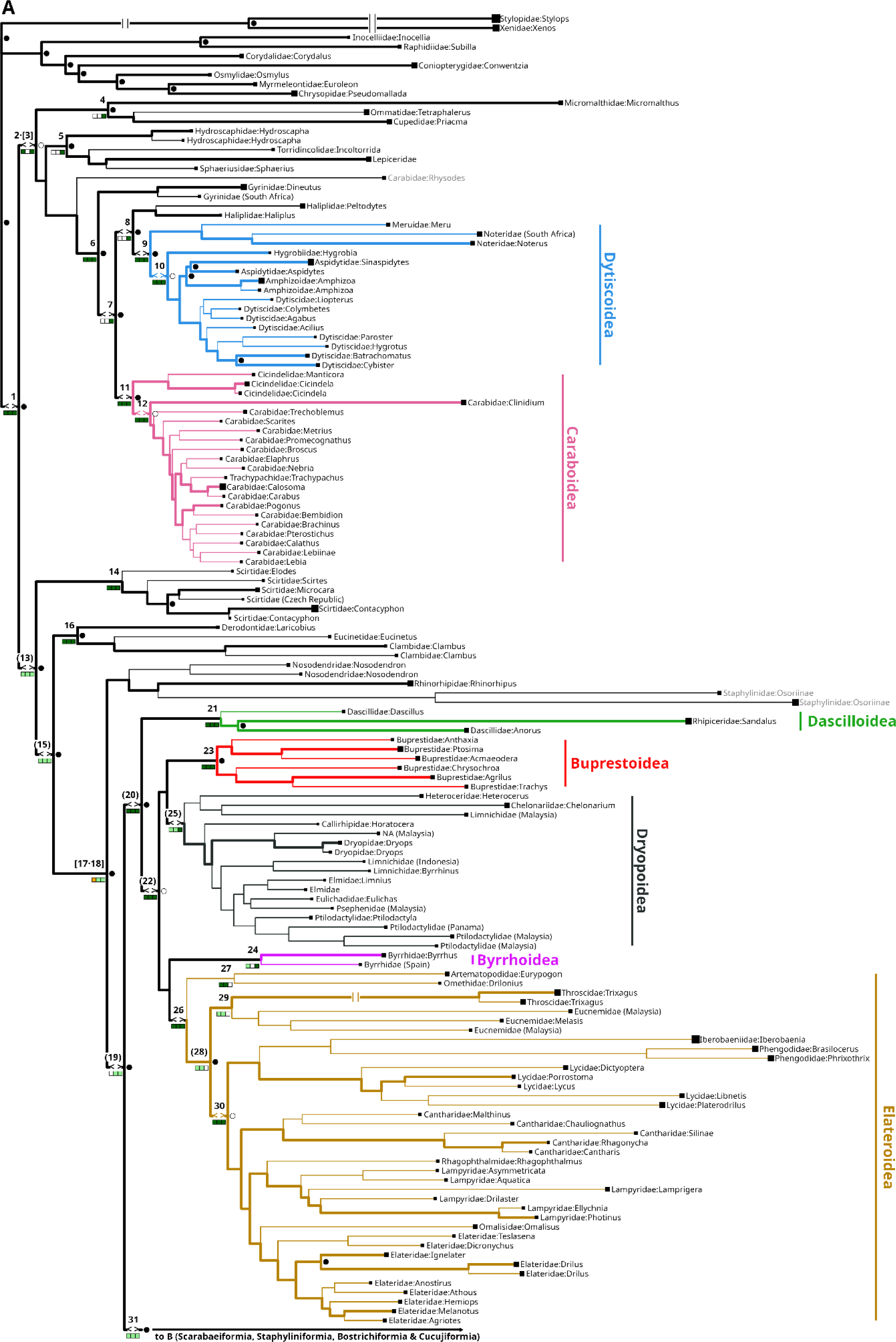
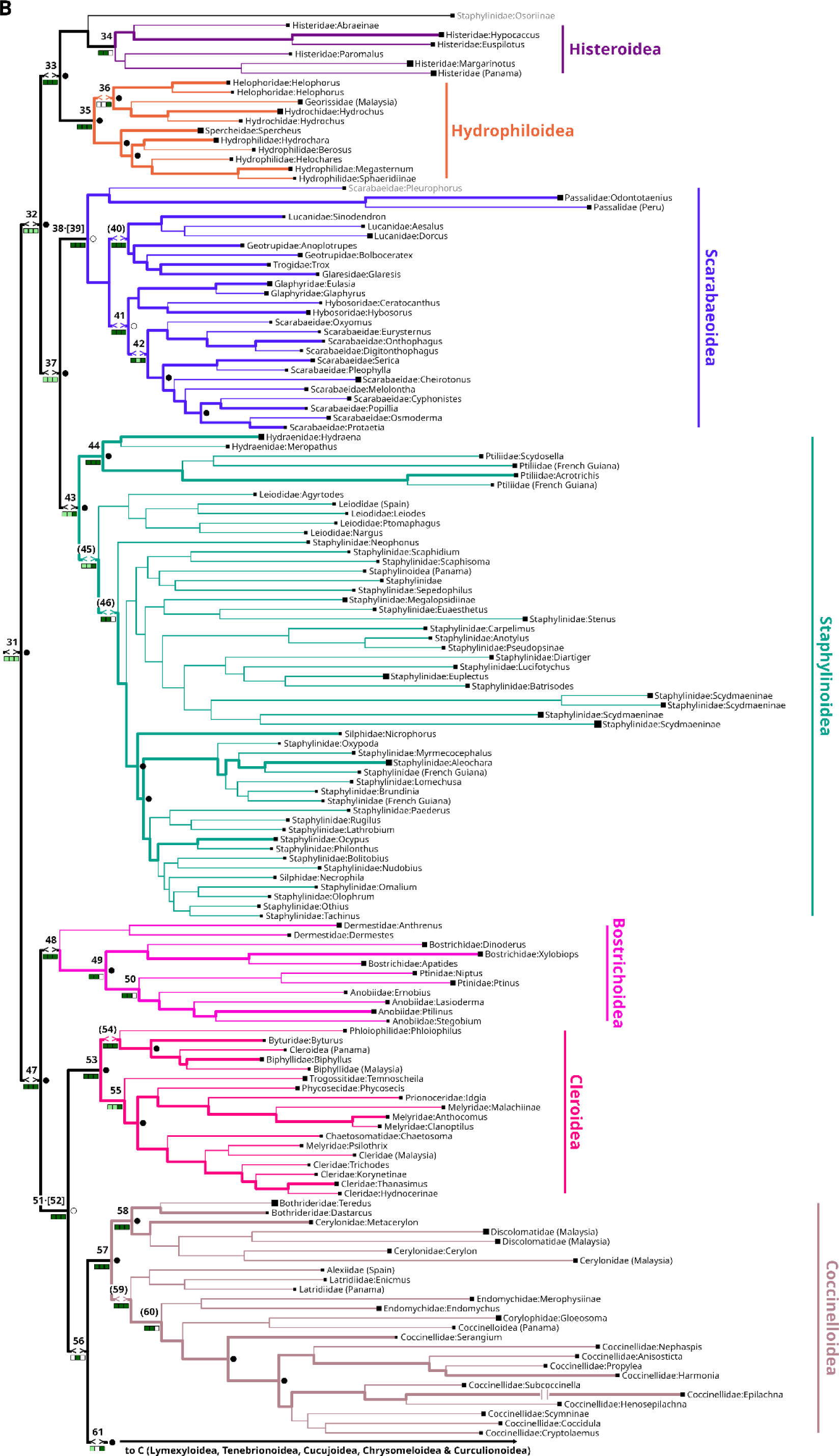
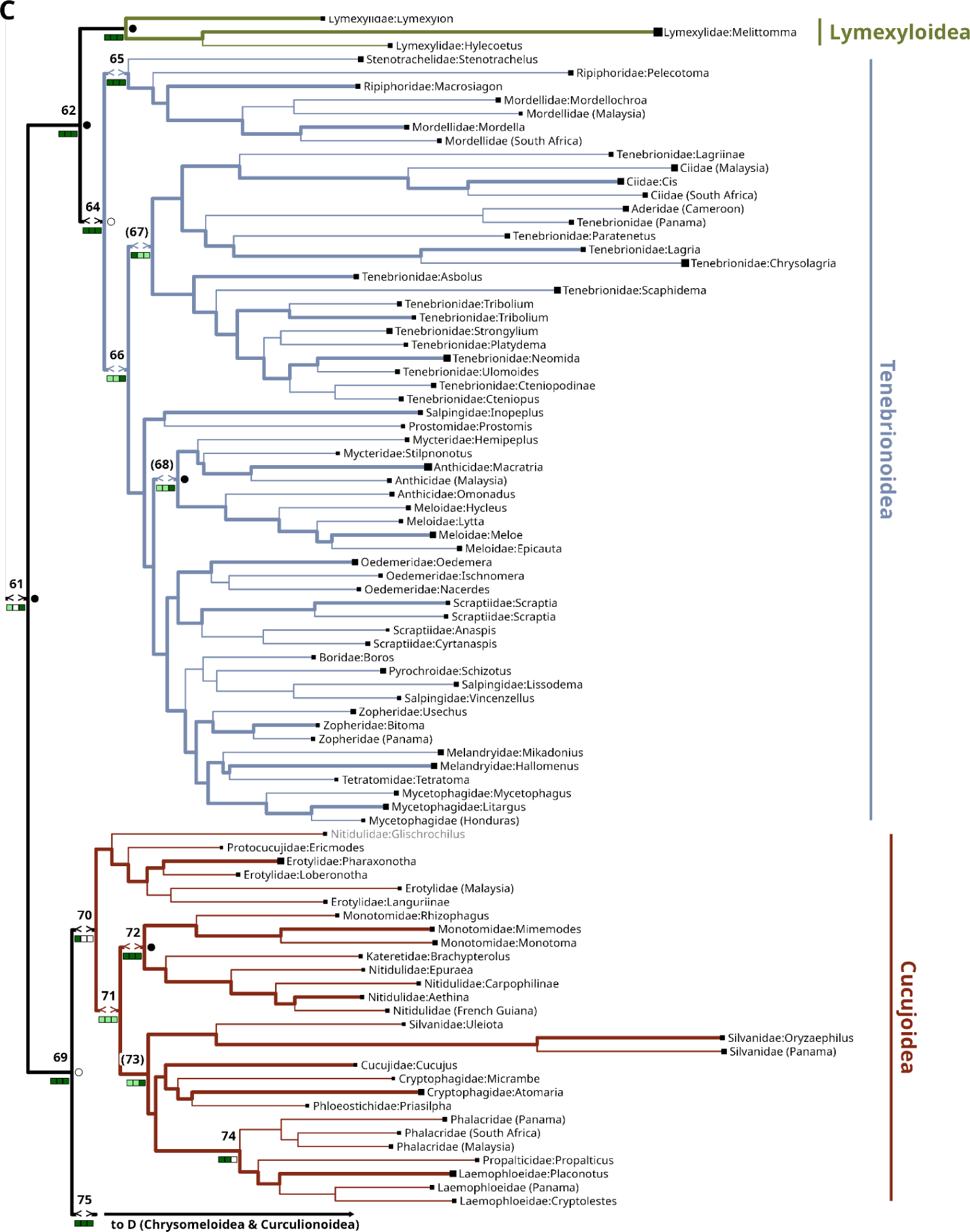
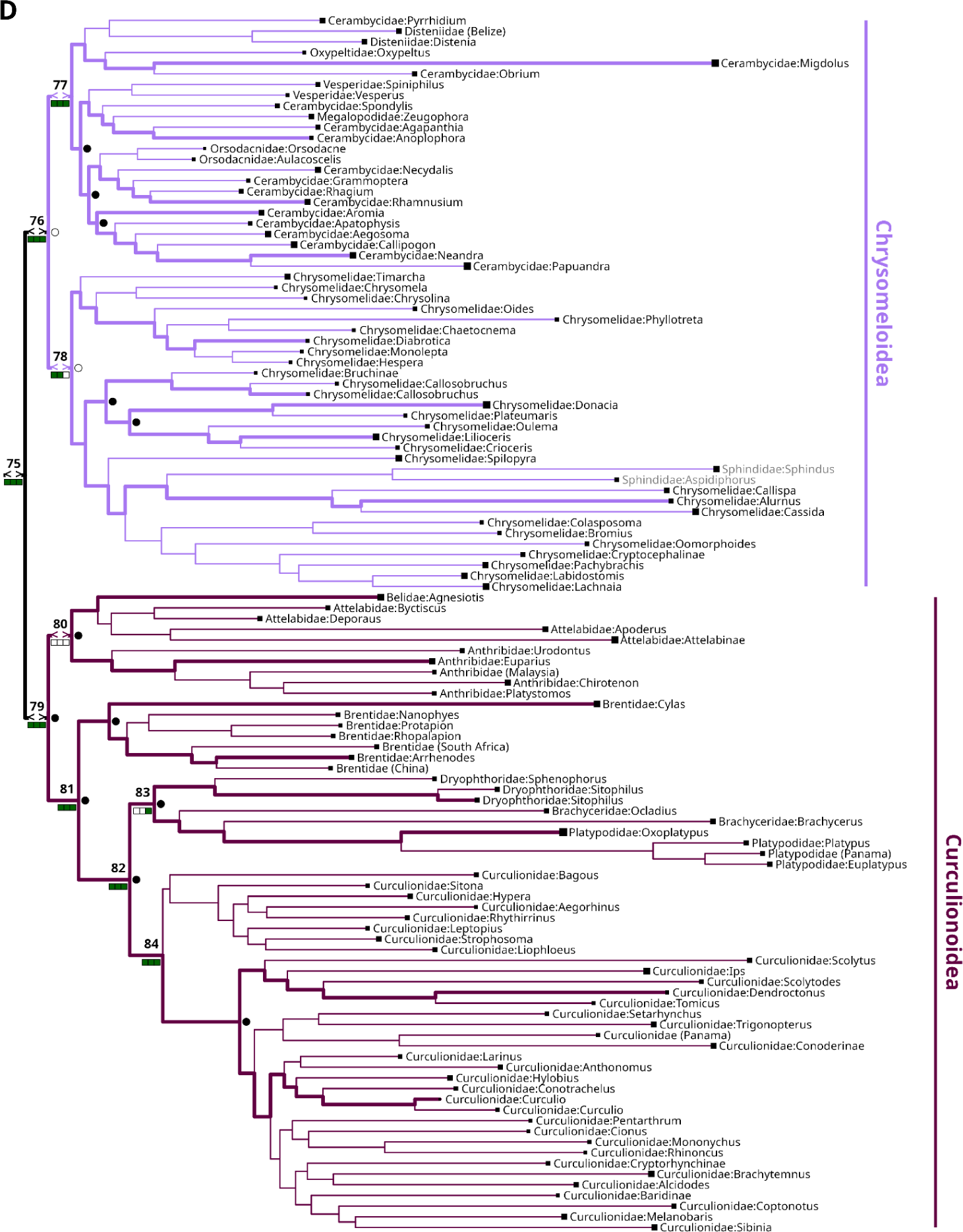
A-D. The mitochondrial amino acid phylogeny of Coleoptera. A tree of 491 mitogenomes generated under a backbone constraint (thick branches). Numbers on the nodes correspond to the MRCA of taxonomic sets derived from the 84 focal clades of the Coleoptera classification at and above the family level, as described in Table 3. Parentheses denote a “monophyletic-incomplete” taxonomic set (i.e. taxa not part of a clade defined in Table 3 are descended from the given node, but these taxa are not part of another group) and brackets denote paraphyletic groups (i.e. taxa that are members of another group in Table 3 are also descended from the node). Circles to the right of internal nodes indicate if a node was in the backbone: circle is filled if it was resolved in the backbone (bifurcation) or empty if it was unresolved (multifurcation). The set of squares adjacent to internal nodes represent the status of this node in three other recent studies (from left to right, Zheng et al. 2018, Cai et al. 2022, McKenna et al. 2021). These are dark green if the group is monophyletic, light green if monophyletic-incomplete, orange if paraphyletic, and white if absent on each tree respectively. Terminal labels give taxonomy: if available, family and genus, otherwise family and subfamily, otherwise the lowest known taxonomy plus the country of origin if known. A few terminals (in grey lettering) are clearly misplaced on this tree and are ignored for the scoring of monophyly. Squares on the tips illustrate taxon-specific compositional heterogeneity, with the side length of each square proportional to the p tsRCFV for that taxon. To improve readability, some short internal branches have been lengthened, denoted by <> in the branch, and some long branches have been shortened, denoted by ||. A single pdf image of the complete tree is available in the supplement, along with a Newick-format file and complete metadata for all terminals.

Within Adephaga, Gyrinidae (Whirligig Beetles) were sister to all other families, which were separated into the traditional terrestrial Geadephaga and aquatic Hydradephaga (Shull et al. 2001). Relationships within Geadephaga were stable, placing Cicindelidae as sister to all others, including Rhysodidae (one outlier) and Carabidae. Within Hydradephaga, various methodologies resulted in different relationships of families in the superfamily Dytiscoidea, with the position of Hygrobiidae shifting, as discussed recently (Vasilikopoulos et al. 2021).

The large suborder Polyphaga at the levels of Series (infraorders) was generally arranged as (Scirtiformia (Elateriformia (Staphyliniformia incl. Scarabaeiformia (Bostrichiformia, Cucujiformia)))). Within the large Cucujiformia comprising approximately half of all described species of Coleoptera, superfamilies were arranged as (Cleroidea (Coccinelloidea ((Lymexyloidea + Tenebionoidea) (Cucujoidea s.str. (Curculionoidea + Chrysomeloidea)))). Within Staphyliniformia internal relationships were ((Hydrophyloidea + Histeroidea) (Staphylinoidea + Scarabaeoidea))). Within Elateriformia we generally observed (Dascilloidea (Elateroidea (Byrrhoidea + Buprestoidea))), although the monophyly of Byrrhoidea was not always recovered. Within these superfamilies, certain subclades were found consistently in our analyses, which also matched the recent phylogenomic studies of McKenna et al. (McKenna et al. 2019), Zhang et al. (Zhang et al. 2018b) and Cai et al. (2022) (Supplementary Figures S3, S4, S5). We thus defined a set of some 84 nodes at family level and above that can be considered a secure scaffold for classification of Coleoptera (Table 3; note this set was arrived at independently from the set of 83 nodes constrained by the nuclear backbone - while these sets overlap, the similarity in the numbers is coincidence). The table also included a few cases where basal relationships were recovered differently in the four studies, including alternative positions of the Cleroidea and Coccinelloidea within Cucujiformia, which require particular attention in future.

**Table 3.**
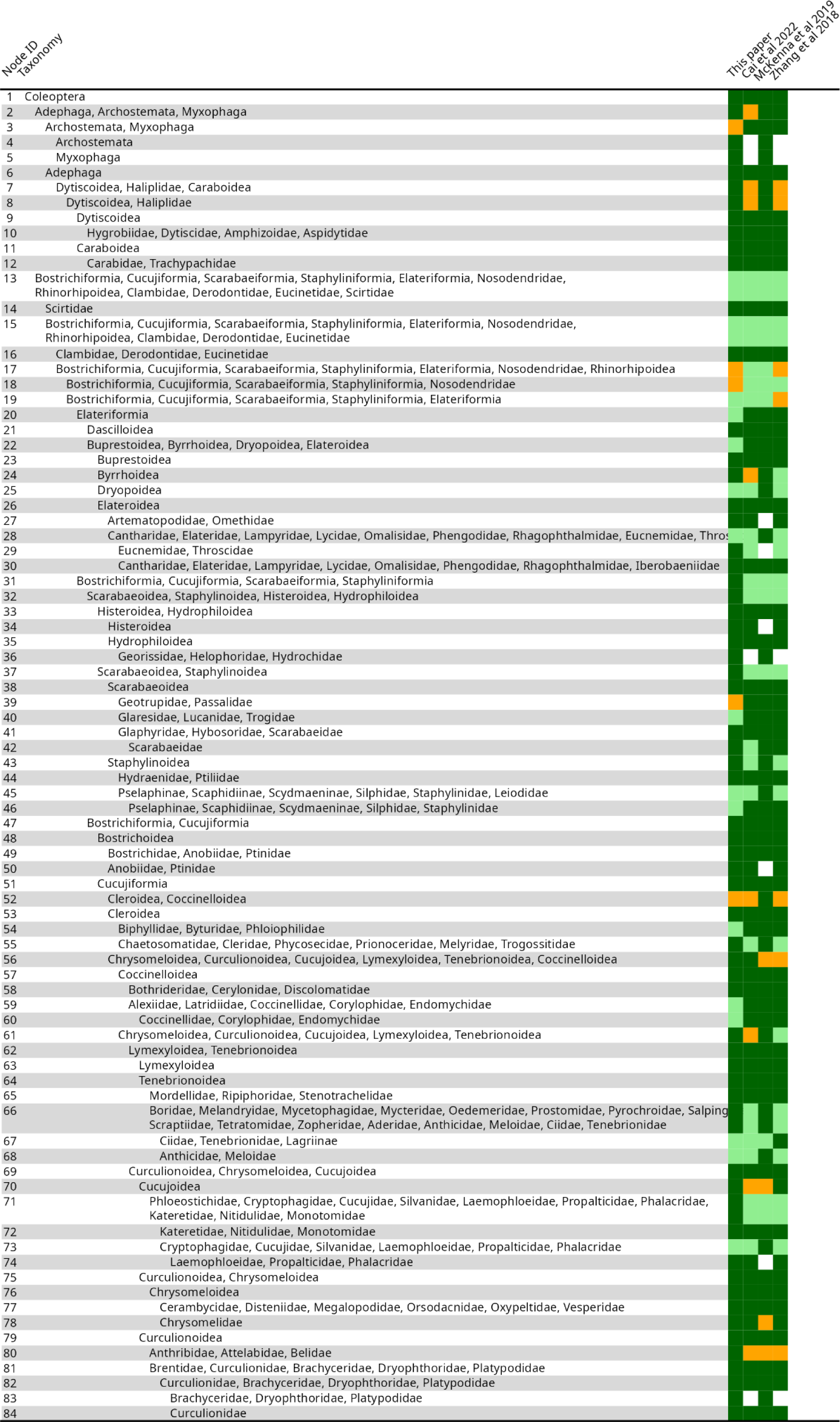
Table of taxonomic comparisons for this and other recent studies. Rows on the table correspond to clades. The taxonomy defines a clade, the Node ID corresponds to the numbers labelling the nodes in the phylogeny on Fig. 6. The coloured boxes correspond to the status of this clade on this and three other recent studies; colours correspond to status: dark green if the group is monophyletic, light green if monophyletic-incomplete, orange if paraphyletic, and white if absent on a given tree (see Fig. 6). Supplementary Material S2 provides Newick files of the four phylogenies with clades mapped to nodes and a csv version of this file.

## Discussion

The rapid growth of phylogenomic sequencing presents unique opportunities and challenges. Mitochondrial genomes play a key role in assembling the Tree-of-Life due to the rapidly increasing taxon density achievable with shotgun sequence data, but bioinformatics and phylogenetic analysis fail to keep up with the speed of data generation. We first developed a much-needed workflow for rapid and accurate annotation of mitochondrial genomes (Cameron 2014), overcoming a major deficiency of existing automated procedures that produce annotations without further tests of alternative, and possibly more plausible assignments of the 5’ and 3’ ends of genes. Second, we developed an approach for integrating the potentially huge numbers of mitogenomes with phylogenetically powerful nuclear orthology sets supporting deep relationships but allowing the more densely sampled mitogenomes to bear on the overall topology. Applied to the Coleoptera, the approach was used to establish >80 nodes that define the basal relationships unlikely to be overturned in future analyses. They can be used as landmarks for further taxon additions and the generation of supertrees. In addition, the taxonomically and biogeographically broadly sampled mitochondrial genome data provide the basis for detailed studies at the family and subfamily levels and allow the robust placement of short barcode and metabarcode sequences.

### Mitogenome workflow

Gene annotation is a key initial step in the processing of mitochondrial genomes. Ambiguity about the precise extent of genes at the 5’ and 3’ ends is due to the surprising density of potential start codons (including non-canonical codons) and stop codons (including partial T and TA codons). Several wrapper scripts are available that connect the various assembly and annotation steps, including the popular MitoZ pipeline (Meng et al. 2019). However, these methods rely on Hidden Markov Model (HMM) profiles for recognition of genes that have limited resolution to establish the start and stop codons and propagate existing annotation biases in the databases. Consequently, genome annotations available at GenBank are rife with errors (Cameron 2023, 2024), and we confirmed this here for numerous submissions of Coleoptera. Our *mitocorrect* tool exploits the largely conserved intergenic distances that define the extent of each gene. Starting with the HMM generated predictions, the method evaluates various alternatives of in-frame annotations defined by different start and stop codons, which each is tested for their deviation from a modal distance to the adjacent genes (within the ranges of distances normally encountered; Suppl. Table 1). The deviations from the modal genome position are then weighted against the deviations from the consensus alignment, for selection of the best annotation. We used this method for a reannotation of published data on Genbank, detecting many inconsistencies of existing annotations (and various frameshifts). The method can greatly speed up the processing of mitogenomes from any source, given a fixed gene order and parameter set. It will be less useful in lineages with more frequent gene order rearrangements, where the context of the adjacent genes is not available. However, alternative gene orders and gene distances can be specified in the custom database, as was done here for the rearrangement of the tRNA-Pro that was known as a clade marker of the Dryopoidea (Timmermans and Vogler 2012), and the method can even serve to detect such rearrangements automatically if the expected neighbour is not present within the custom distance.

The subsequent steps of the workflow address the search for taxonomically defensible trees from the densely sampled mitochondrial markers but avoid the confounding effects of long-branch attraction. The latter presumably is ameliorated by dense, uniform taxon sampling along the deep branches for more accurate inference of the character variation, which we aimed to achieve by the maximum available coverage of the Linnaean taxonomy at family and subfamily levels. However, in many parts of the world the phylogenetic diversity of Coleoptera is too poorly known to use this approach. We therefore added samples from tropical sites representing divergent lineages from a larger database (Creedy, Lee et al., unpublished) and obtained a representation of mitogenomes largely corresponding to the number of species in (super)families of Coleoptera. In the subsequent phylogenetic analysis, standard methods for partitioning and model choice were applied to likelihood searches on both the nucleotide and amino acid datasets, which improved the topologies over unpartitioned searches (Fig. 5). Amino acid coding produced the best trees, and the basal topology was improved under the backbone constraint. However, a few lineages at (sub)family level were placed distantly from the expected positions in all searches, including: Noteridae/Meruidae (Adephaga), Passalidae (Scarabaeoidea), Pselaphinae, Scydmaeninae, Osoriinae (Staphylinoidea), and Sphindidae (Cucujoidea). These groups generally occupied long terminal branches, but did not stand out with regard to compositional heterogeneity (tsRCFV) (Fig. 6), and it remains unclear what features are ultimately causing their aberrant positions. They had to be disregarded when analysing the final trees although they may prove useful for inferring internal relationships of these groups when sampled in more detail.

### Use of the nuclear orthology set

At the deep level, the proposed workflow requires genome sequences or SRA repositories to generate the nuclear orthology set. Availability of genome sequence data in the Insecta is increasing at pace but remains very uneven taxonomically (Feron and Waterhouse 2022). Access to SRA data across major lineages of Coleoptera was greatly enhanced by the work of McKenna et al. (2019) and several other studies. In other situations the primary sequence data may have to be gathered first, although once DNA-grade specimens are at hand, the sequencing methodology is straightforward (Kusy et al. 2018). The nuclear set of some 2,000 loci proved to be powerful in resolving the basal relationships of Coleoptera, but it remained sensitive to model choice and the exact composition of the character matrix, given the trade-off between more genes versus fewer missing characters. We established that the ASTRAL method employing a multispecies coalescent produced poor trees and thus was not considered further; while being fast computationally, and potentially its performance could be improved with weighting of individual gene trees (Zhang and Mirarab 2022), gene concatenation seems more appropriate, as deep coalescence due to ILS is only of limited relevance in the study of basal relationships of a group as ancient as the Coleoptera. Among the remaining 30 tree searches (six matrices x five models), the overall topologies were similar irrespective of the data matrix and model used. Yet, numerous nodes are not universally supported across all treatments, especially near the tips of the tree (Fig. 4). Nodes sensitive to the use of different matrices are also sensitive to the application of different models within a matrix type, suggesting overall weak signal for the affected nodes and possibly insufficient amounts of data. This may be supported by the fact that the smallest (but most complete) *matrix_90* dataset showed comparatively large tree distances from the other trees, which suggests that the size of the matrix is more important than the degree of completeness, in agreement with simulation studies showing that missing characters per se are not as problematic for phylogenetic accuracy as having too few complete characters (Wiens 2003).

In the next step of the protocol, the tree from nuclear data is imposed on the mitogenome tree searches, using a backbone constraint. The strategy for selection of constrained nodes was conservative, requiring maximum (100% bootstrap) support and universal presence across all parameter settings. The great majority of these nodes are also supported by the other major phylogenomic studies (Table 3). The remaining unconstrained nodes generally were closer to the tips (Fig. 4), and therefore were left to be resolved by the mitogenomes, which mostly represent tip level lineages and contribute only a few deep lineages that were not also represented by the nuclear genomes (see the thin branches in Fig. 6).

The link of nuclear and mitochondrial data is via the matrix of mitochondrial sequences required for all included species, including those used for the nuclear genome sequencing. We spent great effort on extracting mitogenomes from the SRA submissions, but most primary nuclear genome data were transcriptomes that did not always assemble into complete error-free contigs. In part, the corresponding mitogenomes could be obtained from Genbank and our unpublished database, and if there was no match on the species level, congenerics or composite sequences were created from multiple Genbank entries of close relatives to represent critical family-level taxa in a few cases (Material and Methods). If relationships in a group are well known, the backbone constraint could be based on an external topology, but again this requires that the specific taxa in the external tree are the same as those used in the mitogenome analysis. This protocol is different from backbone constraints that might be applied based on a classification which would force all members of a higher taxon to adhere to the constraint. We thus avoid the assumption of monophyly of higher taxa, and the large proportion of mitogenomes not represented in the nuclear data are free to ‘find their place’ around the core constraint tree (the fat branches in Fig. 6).

Yet, for many nodes the backbone constraint plays a critical role in resolving basal relationships that are incorrectly recovered using mitogenomes alone. None of the trees constructed under the constraints passed the AU test, indicating a significant divergence from the best tree made from the mitogenomes (Table 2). Yet, the high tRI and tCI values provide validation for the constraint tree topology, especially for the higher levels (suborder, superfamily), which attests to the shortcomings of the mitogenomes on their own. However, the high tCI/tRI for the tree overall also demonstrates the power of mitogenomes if combined with a limited set of nuclear genomes, in particular for the resolution at lower levels (families) where the tRI of constrained and unconstrained searches differ only very little (Table 2). In addition, the mitogenomes, being an independent character system, can arbitrate between contradictory results from nuclear genomes, including the results from previous studies on the Coleoptera phylogeny (Table 3). The backbone constraints are probably the best currently viable option for integrating large nuclear ortholog sets with the more abundant mitogenomes. Tree searches on the nuclear orthology set of ∼2,000 loci are challenging already at the scale of ∼100 terminals used here, and searches on larger taxon numbers in a single matrix including the mitogenomes would make this search very difficult with current algorithms. Anticipating a tree with ∼5,000 mitogenomes (Creedy, Lee, et al., unpublished), backbone constrained tree searches can be achieved with standard phylogenetic software such as IQ-TREE and RAxML. This removes the need for divide-and-conquer methods for tree searches on partial datasets which require the merging of subtrees either by grafting onto a basal phylogeny (Álvarez-Carretero et al. 2022) or tree merging procedures (Zaharias and Warnow 2022) which fuse monophyletic groups that are not subject to further tree searches with respect to each other.

### Classification of Coleoptera

The final objective of this study was to establish an updated framework of the mitogenome phylogeny of the Coleoptera. This was last conducted based on an analysis of ∼250 mitochondrial genomes sampled across the Coleoptera (Timmermans et al. 2016) but now can be extended greatly for a more uniform representation of mitogenomes commensurate with the known species diversity and an overall lower proportion of missing data in the matrix, while basal relationships are supported by the nuclear phylogenomic data. The Coleoptera has been split into nearly 200 families in four suborders and up to 20 superfamilies. Besides the mitogenome study, the two prominent studies of Zhang et al. (2018) and McKenna et al. (McKenna et al. 2019) provide the most complete picture of basal relationships in the order to date. In addition, the former dataset was reanalysed under a CAT model that presumably addresses compositional site-heterogeneity, which also removed potential paralogs (Cai et al. 2022),(Cai et al. 2024). Because these studies are based on very different data types (respectively, PCR of >90 nuclear amplicons; assembly of ∼4,000 loci from short reads; full mitogenomes) and low overlap in taxon choice it has been difficult to synthesise the information, but our survey reveals a set of universally supported deep nodes of Coleoptera. Consequently, we propose a set of ∼80 nodes at family level and above defining the overall frame of the Coleoptera tree against which existing and newly incoming data and analyses can be scored. Most of these nodes are also included in a new classification scheme proposed by Cai et al. (2022) although in some cases the naming of new Series and Superfamilies in this study defines new higher taxa that are not universally supported (see Boudinot et al. 2022). Our set of defined nodes makes the comparison of different studies more precise and quantitative. The current tree also reveals obvious targets for more detailed analyses, especially nearer the tip of the tree, and it defines areas of (mild) disagreement, such as the precise relationships of the infraorders (nodes 17, 18; Fig. 6) and the superfamilies in the Cucujiformia (nodes 52, 56, 61, 70), the question about the monophyly of Cucujoidea (node 70), the position of Byrrhoidea (node 25), and the relationships within Scarabaeoidea and Staphylinoidea. These deep nodes should be confirmed across studies before making changes to the classification. The proposed framework for classification of the Coleoptera provides a solid system into which additional nodes can be added. Defining specific nodes will allow moving away from the current narrative accounts of (dis)agreement that have hampered the comparisons between studies. Our software can place these focal nodes, in whatever way defined, and visualise them on any tree topology. This option also applies to DNA barcode data if aligned to the respective mitogenome region, for a reliable placement of short sequence reads with low distance to a mitogenome sequence.

## Conclusion

As we assemble the animal Tree-of-Life at ever finer levels, mitochondrial genes will continue to play a key role in the study of the most diverse lineages such as the Coleoptera, due to their wide availability and use as barcode markers in biomonitoring. This study addresses the practical implications of using mitochondrial genomes at large scale by providing a streamlined workflow for annotation and alignment. In addition, we provide a methodology for overcoming the well-established shortcomings of mitogenomes in resolving deep nodes by integrating nuclear markers representing the major coleopteran lineages, using partially constrained searches. Until much faster methods for tree searches become available (e.g. Voznica et al. 2022) the use of nuclear genomes in phylogenetics will remain limited to small sets of selected exemplars while for the foreseeable future most of the species diversity will be represented by a few genes only. Mitochondrial genomes are ideal for this purpose and constitute a bridge to the much more abundant (meta)barcoding markers which ultimately need to be incorporated into the complete phylogenetic tree where no other information is available, i.e. so-called ‘dark taxa’ (Hartop et al. 2022). The approach proposed here lays the methodological framework for a seamless integration of phylogenomics, mitogenomics, and DNA barcoding of the Coleoptera.

## Acknowledgements

This work was funded in part by the NHM Biodiversity Initiative (2013-2017), the iBioGen consortium of Horizon 2020, a fellowship by the Chinese Scholarship Council to YD, a Leverhulme International Fellowship IAF-2018-038 to APV, and a philanthropic grant to the NHM. We are grateful to many people supplying specimens and DNA sequences, especially Drs Paula Arribas and Carmelo Andujar (La Laguna, Spain) for sequencing specimens from the NHM Frozen Collection. We thank Peter Foster for computing support.

## Supplementary material

Data available from the Dryad Digital Repository: https://doi.org/10.5061/dryad.zkh1893f4

## Author Contributions

APV and TJC conceived the project. YD carried out nuclear sequence bioinformatics and phylogenetic analysis. TJC wrote the *mitocorrect* tool; TJC and LS carried out mitochondrial sequence bioinformatics. KG undertook an initial version of the mitochondrial phylogenetics; TJC undertook the final version of the mitochondrial phylogenetics. TJC, YD, LS, FZ and KG carried out data analysis and production of figures. KG, APV and TJC wrote the initial draft of the manuscript; all authors contributed to the final draft.

## Supplementary detail on mitoCorrect specifications

The mitocorrect specifications table used to improve annotations for the sequences in this study can be found at https://github.com/tjcreedy/mitocorrect/commit/8852274da53842a368377760bc91a41be0dd9183. Small tweaks have been made since then and we recommend always using the most up-to-date version, as this is derived from the widest possible dataset. The table defines potential start and stop codons in relation to adjacent mitochondrial genes, by specifying the modal distance from (positive values) in the number of nucleotides or overlap (negative values) with the adjacent annotation at the 5’ and 3’ end, respectively. The table also gives the potential start and stop codons, and the maximum range of distances from adjacent genes used to search for potential start and stop codons. For more information, see https://github.com/tjcreedy/mitocorrect#specifications

## Compositional Heterogeneity in the Mitochondrial Dataset

### Methods

Compositional heterogeneity was explored for the three mitochondrial datasets, namely the nucleotide dataset with all three codon positions (NT123) and without the third codon position (NT12), as well as the amino acid dataset (AA). Within each dataset, each partition derived from the partition merging approach (see main text) was considered independently. Within each partition of each dataset, we calculated overall Relative Compositional Frequency Variation (RCFV, Zhong et al 2011) as well as character-specific (csRCFV) and taxon-specific (tsRCFV) components (Kück & Struck, 2014), using custom R functions *rcfv* and *rcfv.partitioned* (https://github.com/tjcreedy/phylostuff/blob/main/phylofuncs.R). For each dataset and partition, we then graphically explored the partitioning of overall RCFV into character-specific and taxon-specific components, and used chi-squared tests to separately test whether the distribution of csRCFV and tsRCFV values across characters and taxa respectively deviated from random chance, i.e. to test whether any characters or taxa appeared to have significantly greater heterogeneity and thus undue influence on the phylogenetic reconstruction.

To summarise tsRCFV across partitions for the amino acid database, we normalised tsRCFV for each taxon within each partition by the total RCFV for that partition, then summed the normalised tsRCFV to generate a single comparable value for each taxon. This value was then plotted on figure X on the main text to explore the potential impact of heterogeneity on the topology of our final tree.

## Results

For partitions within each dataset were renamed as follows:

Amino acid: P1 = ATP6_COX1_COX2_COX3_CYTB_ND3, P2 = ATP8_ND2_ND6, P3 = ND1_ND4_ND4L_ND5

Nucleotide with all three codon positions: P1 = ATP6_1_COX1_1_COX2_1_COX3_1_CYTB_1_CYTB_2, P2 = ATP6_2_COX1_2_COX2_2_COX3_2_ND1_2_ND2_2_ND3_2_ND4_2_ND4L_2_ND5_2_ND6_2, P3 = ATP6_3_ATP8_3_COX1_3_COX2_3_COX3_3_CYTB_3_ND3_3_ND6_3, P4 = ATP8_1_ATP8_2_ND2_1_ND2_3_ND3_1_ND6_1, P5 = ND1_1_ND4_1_ND4L_1_ND5_1, P6 = ND1_3_ND4_3_ND4L_3_ND5_3

Nucleotide without the third codon position: P1 = ATP6_1_COX1_1_COX2_1_COX3_1_CYTB_1_ND2_1_ND3_1_ND6_1, P2 = ATP6_2_ATP8_1_ATP8_2_COX1_2_COX2_2_COX3_2_CYTB_2_ND1_2_ND2_2_ND3_2_ND4_2_ND4 L_2_ND5_2_ND6_2, P3 = ND1_1_ND4_1_ND4L_1_ND5_1

Character-specific compositional heterogeneity was greater on average in the nucleotide datasets, particularly when the third codon position was included (Fig S1). There was moderate variation in csRCFV among partitions, with less variation among partitions in the amino acid dataset.

**Figure S1:**
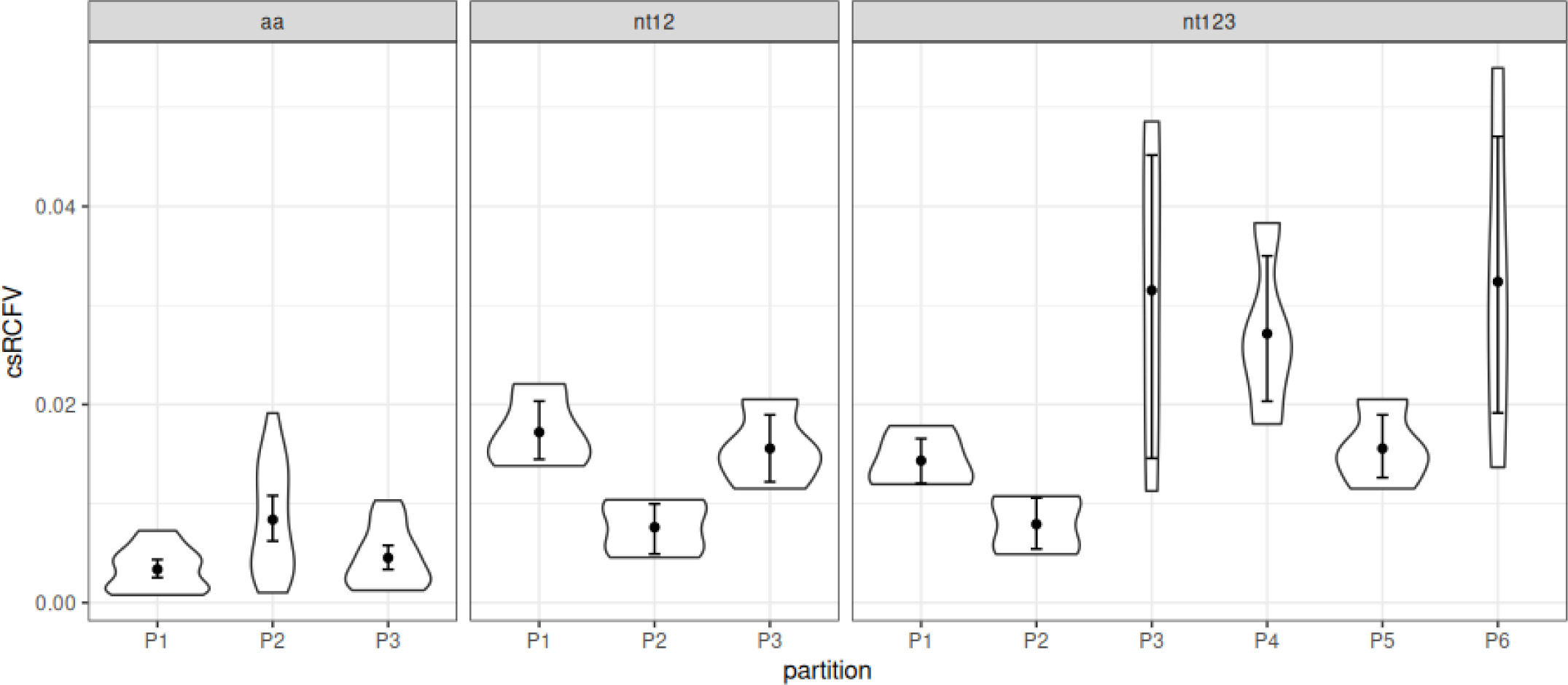
csRCFV values for the three datasets. Plots show the mean csRCFV and bootstrap 95% confidence limits of the mean for each partition within each dataset; violins show the distribution of the data.

While there was some variation in compositional heterogeneity among characters (Fig S2), the chi squared tests of csRCFV value distribution among characters within partitions showed no significance in all cases.

**Figure S2:**
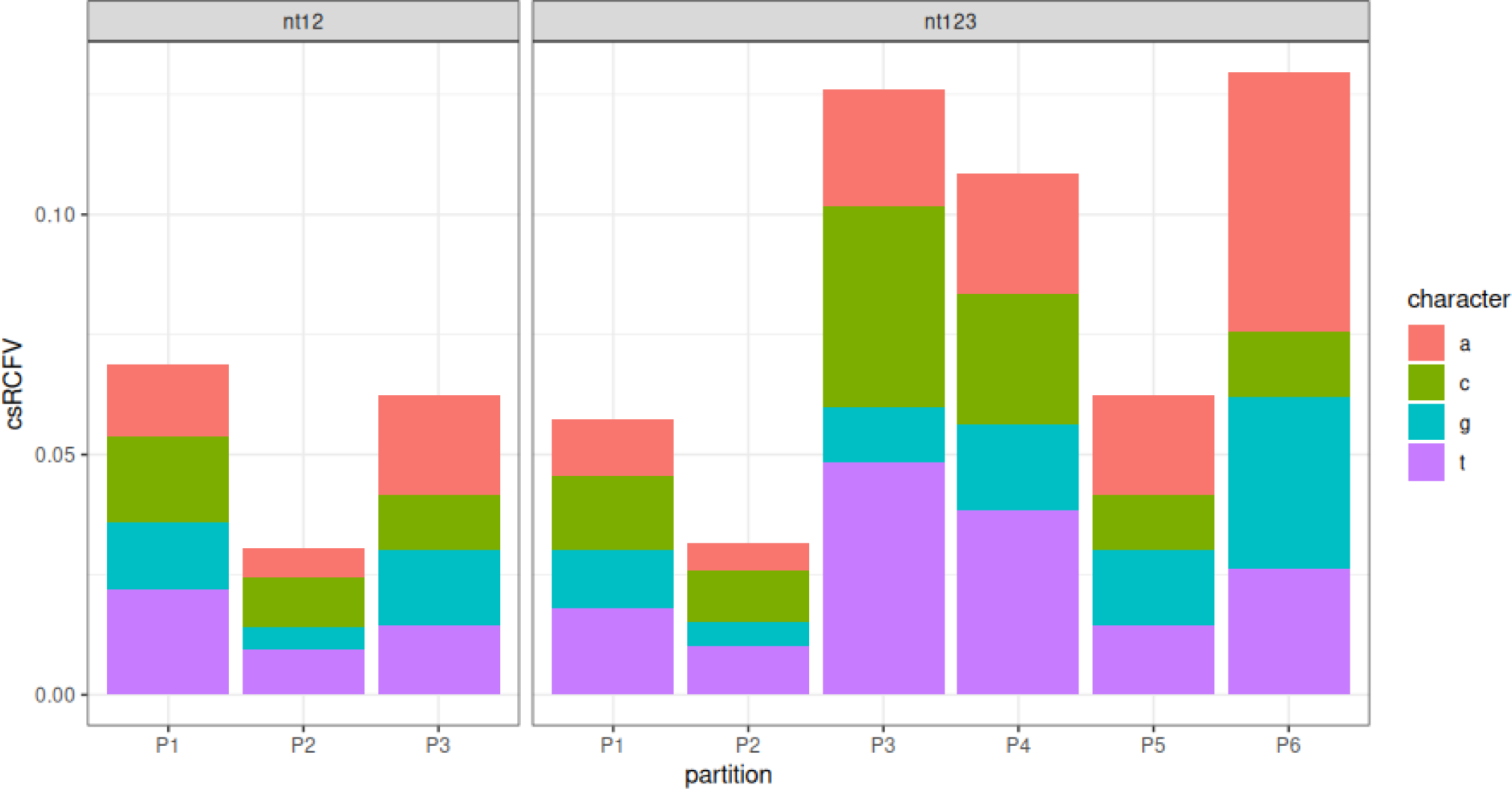
the partitioning of total RCFV (overall bar heights) into character-specific RCFV for the nucleotide datasets. The amino acid equivalent is not shown due to the number of characters.

Taxon-specific compositional heterogeneity was lowest in the nucleotide dataset without the third codon position (Fig S3). While mean tsRCFV was relatively consistent across datasets, the amino acid dataset did have more individual extreme values than the other two datasets. There was minimal difference in compositional heterogeneity between taxa that formed the constrained backbone derived from nuclear data and those found only in the mitochondrial dataset.

**Figure S3:**
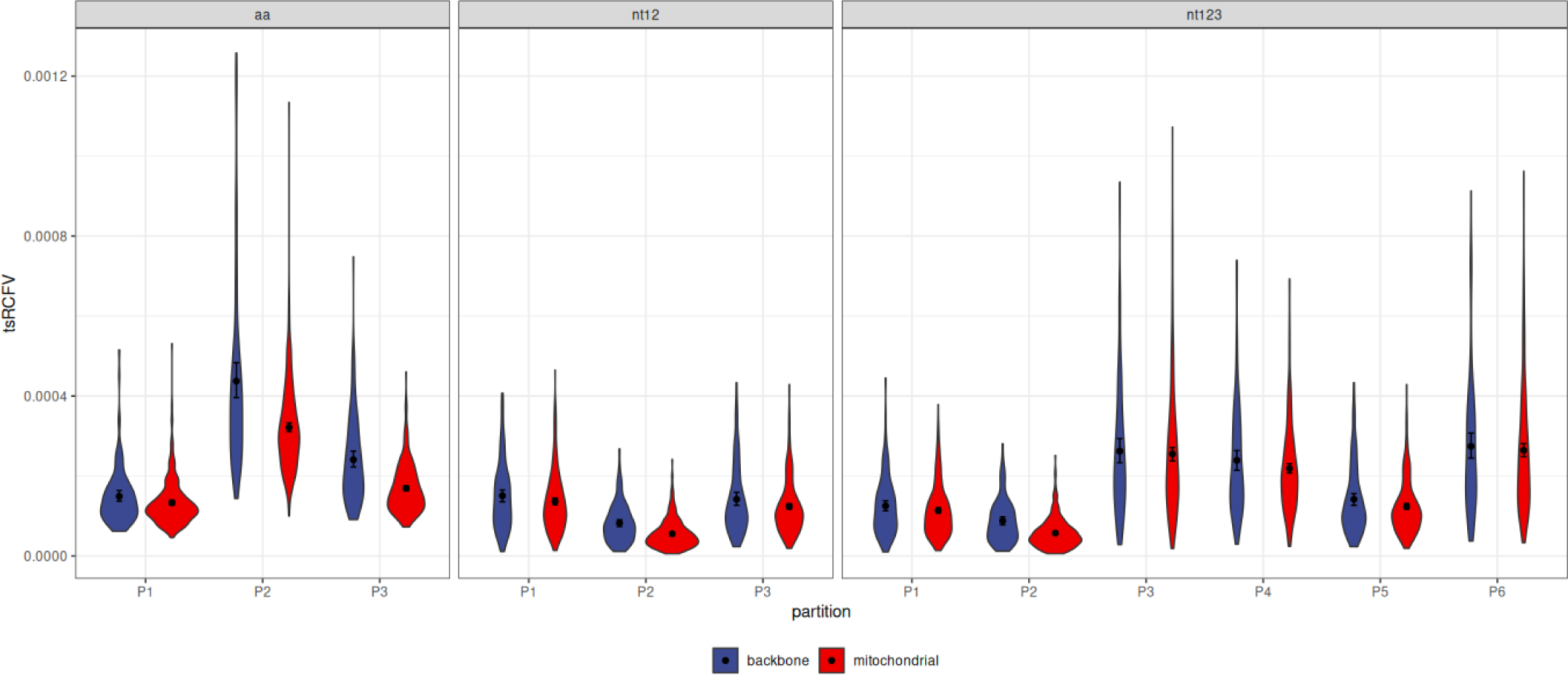
tsRCFV values for the three datasets, split over whether taxa were part of the constrained backbone derived from nuclear data (blue fill), or were found only in the mitochondrial dataset (red fill). Plots show the mean csRCFV and bootstrap 95% confidence limits of the mean for each partition within each dataset; violins show the distribution of the data.

While there was some variation in compositional heterogeneity among taxa, the chi squared tests of csRCFV value distribution among characters within partitions showed no significance in all cases.

## Literature Cited

Adachi J., Hasegawa M. 1996. MOLPHY version 2.3: programs for molecular phylogenetics based on maximum likelihood. Computer Science Monograph.

Álvarez-Carretero S., Tamuri A.U., Battini M., Nascimento F.F., Carlisle E., Asher R.J., Yang Z., Donoghue P.C.J., Dos Reis M. 2022. A species-level timeline of mammal evolution integrating phylogenomic data. Nature. 602:263–267.

Anisimova M., Gil M., Dufayard J.-F., Dessimoz C., Gascuel O. 2011. Survey of branch support methods demonstrates accuracy, power, and robustness of fast likelihood-based approximation schemes. Syst. Biol. 60:685–699.

Blair C. 2023. Organellar DNA continues to provide a rich source of information in the genomics era. Mol. Ecol. 32:2144–2150.

Boudinot B.E., Fikáček M., Lieberman Z.E., Kusy D., Bocak L., Mckenna D.D., Beutel R.G. 2022. Systematic bias and the phylogeny of Coleoptera—A response to Cai et al. (2022) following the responses to Cai et al. (2020). Syst. Entomol. 48:223–232.

Cai C., Tihelka E., Giacomelli M., Lawrence J.F., Ślipiński A., Kundrata R., Yamamoto S., Thayer M.K., Newton A.F., Leschen R.A.B., Gimmel M.L., Lü L., Engel M.S., Bouchard P., Huang D., Pisani D., Donoghue P.C.J. 2022. Integrated phylogenomics and fossil data illuminate the evolution of beetles. R Soc Open Sci. 9:211771.

Cai C.-Y., Tihelka E., Pisani D., Donoghue P.C.J. 2024. Resolving incongruences in insect phylogenomics: A reply to Boudinot et al. (2023). Palaeoentomology. 7:176–183.

Cameron S.L. 2014. Insect mitochondrial genomics: implications for evolution and phylogeny. Annu. Rev. Entomol. 59:95–117.

Cameron S.L. 2023. Mitochondrial phylogenomics of the Australian scribbly-gum moth Ogmograptis (Lepidoptera: Buccaltricidae), and an examination of deep-level relationships within Lepidoptera. Aust. Entomol. 62:449–463.

Cameron S.L. 2024. Insect mitochondrial genomics: A decade of progress. Annu. Rev. Entomol.

Chamberlain S.A., Szöcs E. 2013. taxize: taxonomic search and retrieval in R. F1000Res. 2:191.

Chen S., Zhou Y., Chen Y., Gu J. 2018. fastp: an ultra-fast all-in-one FASTQ preprocessor. Bioinformatics. 34:i884–i890.

Chernomor O., von Haeseler A., Minh B.Q. 2016. Terrace aware data structure for phylogenomic inference from supermatrices. Syst. Biol. 65:997–1008.

Chesters D. 2017. Construction of a species-level Tree of Life for the insects and utility in taxonomic profiling. Syst. Biol. 66:426–439.

Crampton-Platt A., Yu D.W., Zhou X., Vogler A.P. 2016. Mitochondrial metagenomics: letting the genes out of the bottle. Gigascience. 5:15.

Criscuolo A., Gribaldo S. 2010. BMGE (Block Mapping and Gathering with Entropy): a new software for selection of phylogenetic informative regions from multiple sequence alignments. BMC Evol. Biol. 10:210.

Crotty S.M., Minh B.Q., Bean N.G., Holland B.R., Tuke J., Jermiin L.S., Haeseler A.V. 2020. GHOST: Recovering historical signal from heterotachously evolved sequence alignments. Syst. Biol. 69:249–264.

Dayhoff M.O. 1978. A model of evolutionary change in protein. Atlas of protein sequence and structure. 5:345–352.

Desalle R., Schierwater B., Hadrys H. 2017. MtDNA: The small workhorse of evolutionary studies. Front. Biosci. 22:873–887.

Feron R., Waterhouse R.M. 2022. Assessing species coverage and assembly quality of rapidly accumulating sequenced genomes. Gigascience. 11.

Ferrari G., Esselens L., Hart M.L., Janssens S. 2023. Developing the Protocol Infrastructure for DNA Sequencing Natural History Collections. Biodiversity Data Journal. 11:e102317.

Fleming J.F., Struck T.H. 2023. nRCFV: a new, dataset-size-independent metric to quantify compositional heterogeneity in nucleotide and amino acid datasets. BMC Bioinformatics. 24:145.

Fu L., Niu B., Zhu Z., Wu S., Li W. 2012. CD-HIT: accelerated for clustering the next-generation sequencing data. Bioinformatics. 28:3150–3152.

Guindon S., Dufayard J.-F., Lefort V., Anisimova M., Hordijk W., Gascuel O. 2010. New algorithms and methods to estimate maximum-likelihood phylogenies: assessing the performance of PhyML 3.0. Syst. Biol. 59:307–321.

Hartop E., Srivathsan A., Ronquist F., Meier R. 2022. Towards Large-scale Integrative Taxonomy (LIT): resolving the data conundrum for dark taxa. Syst. Biol. 71:1404–142.

Hassanin A. 2006. Phylogeny of Arthropoda inferred from mitochondrial sequences: strategies for limiting the misleading effects of multiple changes in pattern and rates of substitution. Mol. Phylogenet. Evol. 38:100–116.

Hoang D.T., Chernomor O., von Haeseler A., Minh B.Q., Vinh L.S. 2018. UFBoot2: Improving the Ultrafast Bootstrap Approximation. Mol. Biol. Evol. 35:518–522.

Hobern D., Hebert P.D.N. 2019. BIOSCAN-revealing eukaryote diversity, dynamics, and interactions. Biodivers. Inf. Sci. Stand. 3: e37333.

Hunt T., Vogler A.P. 2008. A protocol for large-scale rRNA sequence analysis: towards a detailed phylogeny of Coleoptera. Mol. Phylogenet. Evol. 47:289–301.

Kalyaanamoorthy S., Minh B.Q., Wong T.K.F., von Haeseler A., Jermiin L.S. 2017. ModelFinder: fast model selection for accurate phylogenetic estimates. Nat. Methods. 14:587–589.

Katoh K., Standley D.M. 2013. MAFFT multiple sequence alignment software version 7: improvements in performance and usability. Mol. Biol. Evol. 30:772–780.

Kück P., Longo G.C. 2014. FASconCAT-G: extensive functions for multiple sequence alignment preparations concerning phylogenetic studies. Front. Zool. 11:81.

Kusy D., Motyka M., Andujar C., Bocek M., Masek M., Sklenarova K., Kokas F., Bocakova M., Vogler A.P., Bocak L. 2018. Genome sequencing of Rhinorhipus Lawrence exposes an early branch of the Coleoptera. Front. Zool. 15.

Lanfear R., Calcott B., Ho S.Y.W., Guindon S. 2012. Partitionfinder: combined selection of partitioning schemes and substitution models for phylogenetic analyses. Mol. Biol. Evol. 29:1695–1701.

Lartillot N., Philippe H. 2004. A Bayesian mixture model for across-site heterogeneities in the amino-acid replacement process. Mol. Biol. Evol. 21:1095–1109.

Le S.Q., Gascuel O. 2008. An improved general amino acid replacement matrix. Mol. Biol. Evol. 25:1307–1320.

Le S.Q., Gascuel O. 2010. Accounting for solvent accessibility and secondary structure in protein phylogenetics is clearly beneficial. Syst. Biol. 59:277–287.

Le S.Q., Gascuel O., Lartillot N. 2008. Empirical profile mixture models for phylogenetic reconstruction. Bioinformatics. 24:2317–2323.

Lewin H.A., Richards S., Lieberman Aiden E., Allende M.L., Archibald J.M., Bálint M., Barker K.B., Baumgartner B., Belov K., Bertorelle G., Blaxter M.L., Cai J., Caperello N.D., Carlson K., Castilla-Rubio J.C., Chaw S.-M., Chen L., Childers A.K., Coddington J.A., Conde D.A., Corominas M., Crandall K.A., Crawford A.J., DiPalma F., Durbin R., Ebenezer T.E., Edwards S.V., Fedrigo O., Flicek P., Formenti G., Gibbs R.A., Gilbert M.T.P., Goldstein M.M., Graves J.M., Greely H.T., Grigoriev I.V., Hackett K.J., Hall N., Haussler D., Helgen K.M., Hogg C.J., Isobe S., Jakobsen K.S., Janke A., Jarvis E.D., Johnson W.E., Jones S.J.M., Karlsson E.K., Kersey P.J., Kim J.-H., Kress W.J., Kuraku S., Lawniczak M.K.N., Leebens-Mack J.H., Li X., Lindblad-Toh K., Liu X., Lopez J.V., Marques-Bonet T., Mazard S., Mazet J.A.K., Mazzoni C.J., Myers E.W., O’Neill R.J., Paez S., Park H., Robinson G.E., Roquet C., Ryder O.A., Sabir J.S.M., Shaffer H.B., Shank T.M., Sherkow J.S., Soltis P.S., Tang B., Tedersoo L., Uliano-Silva M., Wang K., Wei X., Wetzer R., Wilson J.L., Xu X., Yang H., Yoder A.D., Zhang G. 2022. The Earth BioGenome Project 2020: Starting the clock. Proc. Natl. Acad. Sci. U. S. A. 119.

Li Z., Linard B., Vogler A.P., Yu D.W., Wang Z. 2023. Phylogenetic diversity only weakly mitigates climate-change-driven biodiversity loss in insect communities. Mol. Ecol. 32:6147–6160.

Luo R., Liu B., Xie Y., Li Z., Huang W., Yuan J., He G., Chen Y., Pan Q., Liu Y., Tang J., Wu G., Zhang H., Shi Y., Liu Y., Yu C., Wang B., Lu Y., Han C., Cheung D.W., Yiu S.-M., Peng S., Xiaoqian Z., Liu G., Liao X., Li Y., Yang H., Wang J., Lam T.-W., Wang J. 2012. SOAPdenovo2: an empirically improved memory-efficient short-read de novo assembler. Gigascience. 1:18.

Manni M., Berkeley M.R., Seppey M., Simão F.A., Zdobnov E.M. 2021. BUSCO update: Novel and streamlined workflows along with broader and deeper phylogenetic coverage for scoring of eukaryotic, prokaryotic, and viral genomes. Mol. Biol. Evol. 38:4647–4654.

McKenna D.D., Shin S., Ahrens D., Balke M., Beza-Beza C., Clarke D.J., Donath A., Escalona H.E., Friedrich F., Letsch H., Liu S., Maddison D., Mayer C., Misof B., Murin P.J., Niehuis O., Peters R.S., Podsiadlowski L., Pohl H., Scully E.D., Yan E.V., Zhou X., Ślipiński A., Beutel R.G. 2019. The evolution and genomic basis of beetle diversity. Proc. Natl. Acad. Sci. U. S. A. 116:24729–24737.

Meng G., Li Y., Yang C., Liu S. 2019. MitoZ: a toolkit for animal mitochondrial genome assembly, annotation and visualization. Nucleic Acids Res. 47:e63.

Minh B.Q., Lanfear R., Trifinopoulos J., Schrempf D., Schmidt H.A. 2021. IQ-TREE version 2.1. 2: Tutorials and Manual Phylogenomic software by maximum likelihood.

Misof B., Liu S., Meusemann K., Peters R.S., Donath A., Mayer C., Frandsen P.B., Ware J., Flouri T., Beutel R.G., Niehuis O., Petersen M., Izquierdo-Carrasco F., Wappler T., Rust J., Aberer A.J., Aspöck U., Aspöck H., Bartel D., Blanke A., Berger S., Böhm A., Buckley T.R., Calcott B., Chen J., Friedrich F., Fukui M., Fujita M., Greve C., Grobe P., Gu S., Huang Y., Jermiin L.S., Kawahara A.Y., Krogmann L., Kubiak M., Lanfear R., Letsch H., Li Y., Li Z., Li J., Lu H., Machida R., Mashimo Y., Kapli P., McKenna D.D., Meng G., Nakagaki Y., Navarrete-Heredia J.L., Ott M., Ou Y., Pass G., Podsiadlowski L., Pohl H., von Reumont B.M., Schütte K., Sekiya K., Shimizu S., Slipinski A., Stamatakis A., Song W., Su X., Szucsich N.U., Tan M., Tan X., Tang M., Tang J., Timelthaler G., Tomizuka S., Trautwein M., Tong X., Uchifune T., Walzl M.G., Wiegmann B.M., Wilbrandt J., Wipfler B., Wong T.K.F., Wu Q., Wu G., Xie Y., Yang S., Yang Q., Yeates D.K., Yoshizawa K., Zhang Q., Zhang R., Zhang W., Zhang Y., Zhao J., Zhou C., Zhou L., Ziesmann T., Zou S., Li Y., Xu X., Zhang Y., Yang H., Wang J., Wang J., Kjer K.M., Zhou X. 2014. Phylogenomics resolves the timing and pattern of insect evolution. Science. 346:763–767.

Nylander J. 2010. catfasta2phyml.

Paradis E., Schliep K. 2019. ape 5.0: an environment for modern phylogenetics and evolutionary analyses in R. Bioinformatics. 35:526–528.

Robinson D.F., Foulds L.R. 1981. Comparison of phylogenetic trees. Math. Biosci. 53:131–147.

Rota-Stabelli O., Yang Z., Telford M.J. 2009. MtZoa: a general mitochondrial amino acid substitutions model for animal evolutionary studies. Mol. Phylogenet. Evol. 52:268–272.

Sayyari E., Mirarab S. 2016. Fast coalescent-based computation of local branch support from quartet frequencies. Mol. Biol. Evol. 33:1654–1668.

Schliep K.P. 2011. phangorn: phylogenetic analysis in R. Bioinformatics. 27:592–593.

Shull V.L., Vogler A.P., Baker, Maddison D.R., Hammond P.M. 2001. Sequence alignment of 18S ribosomal RNA and the basal relationships of adephagan beetles: evidence for monophyly of aquatic families and the placement of Trachypachidae. Syst. Biol. 50:945–969.

Team R.C. 2013. R: A language and environment for statistical computing. R Foundation for Statistical Computing. (No Title).

Timmermans M.J.T.N., Barton C., Haran J., Ahrens D., Culverwell C.L., Ollikainen A., Dodsworth S., Foster P.G., Bocak L., Vogler A.P. 2016. Family-level sampling of mitochondrial genomes in Coleoptera: Compositional heterogeneity and phylogenetics. Genome Biol. Evol. 8:161–175.

Timmermans M.J.T.N., Vogler A.P. 2012. Phylogenetically informative rearrangements in mitochondrial genomes of Coleoptera, and monophyly of aquatic elateriform beetles (Dryopoidea). Mol. Phylogenet. Evol. 63:299–304.

Toups B.S., Thomson R.C., Brown J.M. 2024. Complex models of sequence evolution improve fit, but not gene tree discordance, for tetrapod mitogenomes. Syst. Biol.

Vasilikopoulos A., Balke M., Kukowka S., Pflug J.M., Martin S., Meusemann K., Hendrich L., Mayer C., Maddison D.R., Niehuis O., Beutel R.G., Misof B. 2021. Phylogenomic analyses clarify the pattern of evolution of Adephaga (Coleoptera) and highlight phylogenetic artefacts due to model misspecification and excessive data trimming. Syst. Entomol. 46:991–1018.

Voznica J., Zhukova A., Boskova V., Saulnier E., Lemoine F., Moslonka-Lefebvre M., Gascuel O. 2022. Deep learning from phylogenies to uncover the epidemiological dynamics of outbreaks. Nat. Commun. 13:3896.

Wang H.-C., Minh B.Q., Susko E., Roger A.J. 2018. Modeling site heterogeneity with posterior mean site frequency profiles accelerates accurate phylogenomic estimation. Syst. Biol. 67:216–235.

Waterhouse R.M., Seppey M., Simão F.A., Manni M., Ioannidis P., Klioutchnikov G., Kriventseva E.V., Zdobnov E.M. 2018. BUSCO applications from quality assessments to gene prediction and phylogenomics. Mol. Biol. Evol. 35:543–548.

Waterhouse R.M., Seppey M., Simão F.A., Zdobnov E.M. 2019. Using BUSCO to assess insect genomic resources. Methods Mol. Biol. 1858:59–74.

Wiens J.J. 2003. Missing data, incomplete taxa, and phylogenetic accuracy. Syst. Biol. 52:528–538.

Xie Y., Wu G., Tang J., Luo R., Patterson J., Liu S., Huang W., He G., Gu S., Li S., Zhou X., Lam T.-W., Li Y., Xu X., Wong G.K.-S., Wang J. 2014. SOAPdenovo-Trans: de novo transcriptome assembly with short RNA-Seq reads. Bioinformatics. 30:1660–1666.

Zaharias P., Warnow T. 2022. Recent progress on methods for estimating and updating large phylogenies. Philos. Trans. R. Soc. Lond. B Biol. Sci. 377:20210244.

Zhang C., Mirarab S. 2022. Weighting by gene tree uncertainty improves accuracy of quartet-based species trees. Mol. Biol. Evol. 39.

Zhang C., Rabiee M., Sayyari E., Mirarab S. 2018a. ASTRAL-III: polynomial time species tree reconstruction from partially resolved gene trees. BMC Bioinformatics. 19:153.

Zhang G., Li C., Li Q., Li B., Larkin D.M., Lee C., Storz J.F., Antunes A., Greenwold M.J., Meredith R.W., Ödeen A., Cui J., Zhou Q., Xu L., Pan H., Wang Z., Jin L., Zhang P., Hu H., Yang W., Hu J., Xiao J., Yang Z., Liu Y., Xie Q., Yu H., Lian J., Wen P., Zhang F., Li H., Zeng Y., Xiong Z., Liu S., Zhou L., Huang Z., An N., Wang J., Zheng Q., Xiong Y., Wang G., Wang B., Wang J., Fan Y., da Fonseca R.R., Alfaro-Núñez A., Schubert M., Orlando L., Mourier T., Howard J.T., Ganapathy G., Pfenning A., Whitney O., Rivas M.V., Hara E., Smith J., Farré M., Narayan J., Slavov G., Romanov M.N., Borges R., Machado J.P., Khan I., Springer M.S., Gatesy J., Hoffmann F.G., Opazo J.C., Håstad O., Sawyer R.H., Kim H., Kim K.-W., Kim H.J., Cho S., Li N., Huang Y., Bruford M.W., Zhan X., Dixon A., Bertelsen M.F., Derryberry E., Warren W., Wilson R.K., Li S., Ray D.A., Green R.E., O’Brien S.J., Griffin D., Johnson W.E., Haussler D., Ryder O.A., Willerslev E., Graves G.R., Alström P., Fjeldså J., Mindell D.P., Edwards S.V., Braun E.L., Rahbek C., Burt D.W., Houde P., Zhang Y., Yang H., Wang J., Avian Genome Consortium, Jarvis E.D., Gilbert M.T.P., Wang J. 2014. Comparative genomics reveals insights into avian genome evolution and adaptation. Science. 346:1311–1320.

Zhang S.-Q., Che L.-H., Li Y., Dan Liang, Pang H., Ślipiński A., Zhang P. 2018b. Evolutionary history of Coleoptera revealed by extensive sampling of genes and species. Nat. Commun. 9:205.

Zhong M., Hansen B., Nesnidal M., Golombek A., Halanych K.M., Struck T.H. 2011. Detecting the symplesiomorphy trap: a multigene phylogenetic analysis of terebelliform annelids. BMC Evol. Biol. 11:369.

## Literature cited

Zhong, M., Hansen, B., Nesnidal, M., et al. Detecting the symplesiomorphy trap: a multigene phylogenetic analysis of terebelliform annelids. BMC Evol Biol 11, 369 (2011). 10.1186/1471-2148-11-369

Kück, P., Struck, T. H.. BaCoCa – A heuristic software tool for the parallel assessment of sequence biases in hundreds of gene and taxon partitions. Molecular Phylogenetics and Evolution 70 (2014). 10.1016/j.ympev.2013.09.011

